# ResPAN: a powerful batch correction model for scRNA-seq data through residual adversarial networks

**DOI:** 10.1101/2021.11.08.467781

**Authors:** Yuge Wang, Tianyu Liu, Hongyu Zhao

## Abstract

**Motivation:** With the advancement of technology, we can generate and access large-scale, high dimensional and diverse genomics data, especially through single-cell RNA sequencing (scRNA-seq). However, integrative downstream analysis from multiple scRNA-seq datasets remains challenging due to batch effects.

**Results:** In this paper, we propose a light-structured deep learning framework called ResPAN for scRNA-seq data integration. ResPAN is based on Wasserstein Generative Adversarial Network (WGAN) combined with random walk mutual nearest neighbor pairing and fully skip-connected autoencoders to reduce the differences among batches. We also discuss the limitations of existing methods and demonstrate the advantages of our model over seven other methods through extensive benchmarking studies on both simulated data under various scenarios and real datasets across different scales. Our model achieves leading performance on both batch correction and biological information conservation and maintains scalable to datasets with over half a million cells.

**Availability:** An open-source implementation of ResPAN and scripts to reproduce the results can be downloaded from: https://github.com/AprilYuge/ResPAN.

## 1 Introduction

The advancement of single-cell sequencing technologies and the vast amounts of data generated has contributed significantly to biological research. One major challenge in single-cell data analysis is the removal of batch effect, which refers to the differences in the distributions of the amount of single-cell RNA sequences for cells from the same tissue. This problem is present in essentially all single-cell studies (Papalexi and Satija, 2018; Pijuan-Sala *et al*., 2018; Suvà and Tirosh, 2019). Batch effects can result from various experimental factors, including different protocols, instruments, operators, and pre-processing methods (Leek *et al*., 2010). The removal of batch effect is not only important for us to identify true biological signals but also to facilitate integrative analyses across studies.

Two main approaches have been proposed to reduce batch effects in published work (Luecken *et al*., 2021). The first one is to assume that data from different batches follow the same distribution so that we can estimate the parameters in the distribution and remove the batch effects through statistical modeling. The second one is to select one batch as the reference batch and others as query batches and then get the batch-effect-corrected data by constructing a mapping from the query batches to the reference batch. Representative methods for the first strategy include Limma (Ritchie *et al*., 2015), Combat (Johnson *et al*., 2007), Liger (Welch *et al*., 2019), and scVI (Lopez *et al*., 2018). Representative methods for the second strategy include MMD-ResNet (Shaham *et al*., 2017), Mutual Nearest Neighbor (MNN) (Haghverdi *et al*., 2018), Seurat v4 (Stuart *et al*., 2019), Batch Balanced kNN (BBKNN) (Polański *et al*., 2020), iMAP (Wang *et al*., 2021a), and Harmony (Korsunsky *et al*., 2019).

Even with the many methods available, there remain many challenges for batch correction. The most crucial point is to find a way to perfectly mix cells from the same cell type across different batches and preserve batch-specific biological information (e.g. batch-specific cell types (Wang *et al*., 2021a)). Current methods have different focuses on the above two points, and may lead to poor results after batch effect removal (see Discussion). Therefore, how to design an efficient method to achieve a reasonable trade-off between batch mixing and biological information conservation remains a challenge in practice.

Moreover, the variety of single-cell datasets and the volume of data are increasing, challenging the assumptions made in some of the batch effect removal methods. For example, some statistical and computational methods are built based on linear models (Limma and Liger) or assume that single-cell data follow a specific distribution (Combat and scVI) with parameters associated with the distribution need to be estimated (e.g. mean and variance/dispersion). Moreover, some models represent batch effects with a single batch variable and aim to remove the effect of the batch variable from the estimated distributions (Limma and scVI). However, these assumptions may not hold for many single-cell datasets, because the sources of batch effects can be complicated and entangled with true biological information, so it is not practical to summarize them in one variable or in a specified distribution family. In addition, the accumulating data make it more challenging for these algorithms to run with manageable time complexity. As a result, traditional models may not be the best candidates for batch effect removal for large and complex single-cell data. Therefore, researchers have been adopting machine learning models based on approximation theory, where there is more flexibility in the distribution of input data (Hornik *et al*., 1989), such as generative neural networks implemented in MMD-ResNet, scVI and iMAP. However, there are issues with these three methods. For MMD-ResNet, the training data are not properly pre-processed, often generating over or under corrected data (Tran *et al*., 2020). For scVI, as mentioned above, it represents batch effects as a single variable, making it challenging to generalize well to some data. For iMAP, although it carefully considers data selection through MNN pairs, it adopts a cumbersome network design, and we found that its performance varied under different random seeds, which will be discussed later in this paper. Therefore, the existing machine learning-based correction models are poorly generalized, with varying results across different datasets, due to inappropriate data preprocessing, model assumption and the choice of model structure.

To address the limitations of the existing methods, we propose ResPAN: **Res**idual autoencoder and mutual nearest neighbor **P**aring guided **A**dversarial **N**etwork for batch effect removal. This model has a lighter structure, a relatively shorter training time (through both hardware and software acceleration), and better stability and generalization. These benefits are demonstrated through our benchmarking results.

## 2 Materials and methods

### 2.1 Overview of ResPAN

The workflow of ResPAN has three key steps: generation of training data, adversarial training of the neural network, and generation of corrected data without batch effect (Fig. 1).

**Figure 1:**
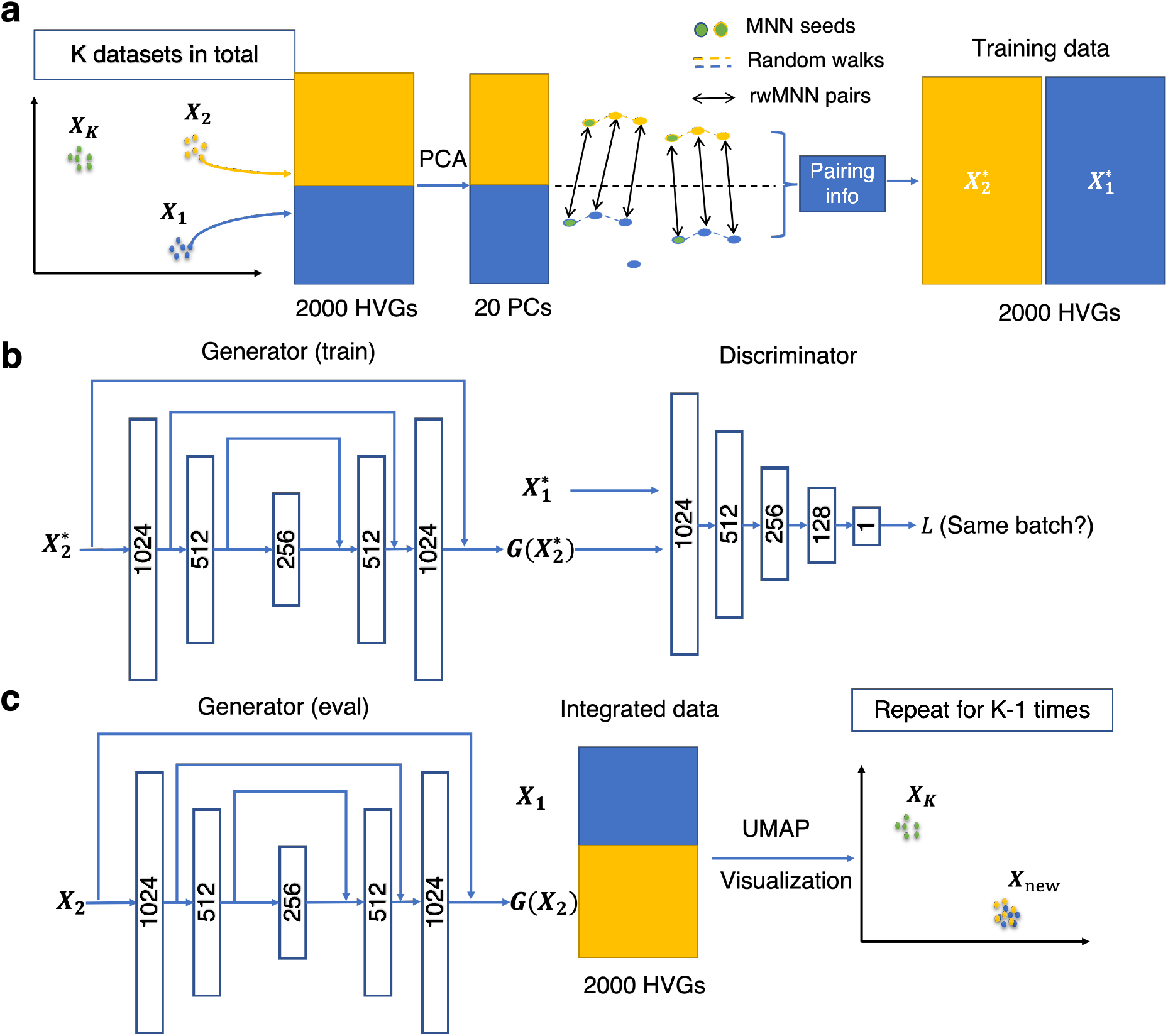
Workflow of ResPAN. (a) Generation of training data based on random walk mutual nearest neighbors (rwMNN) pairs. Pairs are generated in the top 20 PC space, but the training data are in the gene space using all cells contained in the rwMNN pairs. (b) Training process. We utilize the adversarial training strategy to optimize our model. The generator is a fully skip connected version of residual autoencoder. (c) Data integration and visualization. The results received from the generator are utilized for downstream analysis. Notations: *X*_1_: the selected reference batch; *X*_2_: the first query batch; 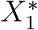: the reference training data; 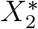: the query training data; *G:* the generator.

In the first step (Fig. 1a), the top 2,000 highly variable genes (HVGs) are selected across all batches as common features for data integration. Inspired by the idea of using mutual nearest neighbors (MNNs) for batch correction (Haghverdi *et al*., 2018; Stuart *et al*., 2019), ResPAN selects training data by finding MNN pairs using cosine similarity between the reference data and the query data on the top 20 principal components (PCs). Since the purpose of GAN is to generate data from one distribution to another and we do not want to lose batch-specific information during training, it is important to make the training data from two batches have similar distributions (e.g. cell type composition) by implementing MNN pairing. However, the number of MNN pairs is too small to be utilized for training our model effectively. Therefore, after finding MNN pairs, ResPAN follows the recommendation in iMAP to augment the training data through random walks. Random walk MNN (rwMNN) pairs are generated by using each MNN pair as the starting seed, and they successively search through the k nearest neighbors (kNNs) of the current cell within the same batch. We found that the mismatch rates of MNN pairs and rwMNN pairs across different datasets are comparable (Supplementary Table S1). Cells selected as rwMNN pairs are used as training data, and note that the same cell may be selected more than once by design. Details of rwMNN pairing can be found in Supplementary Note S1.1.

In the second step (Fig. 1b), after generating the training data, we utilize a modified version of WGAN (Arjovsky *et al*., 2017) with gradient penalty (Gulrajani *et al*., 2017) (WGAN-GP) to generate a mapping function that can transfer the query data so that their distribution is similar to the reference data. WGAN-GP is composed of a generator (*G*) and a discriminator (*D*). The former model is an autoencoder used to generate data that follow a new distribution from the query data, while the latter model is used to discriminate whether the corrected query data have a distribution similar to the reference data. Different from the traditional WGAN-GP, as used in iMAP, we use a fully skip connected residual autoencoder. Skip connection is used for each symmetric pair of layers in the encoder and the decoder. The loss function of WGAN-GP is:

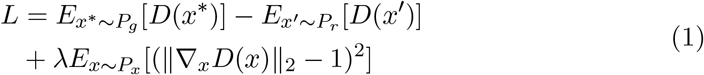

where *P_g_* is the distribution of the data generated from the generator, *P_r_* is the distribution of the reference data, and *P_x_* is the distribution of the data interpolated using the generated and the reference data. The first two terms represent the Wasserstein Distance and the third term is the gradient penalty used to enforce Lipschitz continuity (Gulrajani *et al*., 2017). When training *D*, we aim to minimize the loss function; while when training *G*, we aim to maximize it. Such adversarial training strategy is adopted to train *D* and *G* alternately and finally achieve the Nash equilibrium between them (Heusel *et al*., 2017). More details about the neural network structure of ResPAN and its training process can be found in Supplementary Note S1.2.

In the last step (Fig. 1c), we use the trained generator to map all cells in the query batch to the reference batch and combine it with the reference batch to get an integrated dataset. When there are two batches, we only need to perform the aforementioned process once. When there are *K*(*K* > 2) batches, we utilize the sequential training strategy by running the training data generation and the network training process for *K* – 1 times. After each round, one query batch is integrated into the entire reference batch, making the reference batch larger and larger. This strategy is adopted because different query batches may have different distributions. The integrated data is used for downstream analysis, including data visualization and method benchmarking.

### 2.2 Dataset information

For the simulation study, we generated synthetic scRNA-seq data using Splatter (Zappia *et al*., 2017), which is based on negative binomial distributions through a hierarchical Gamma-Poisson model. Parameters were the same as those used in Kotliar et al. (Kotliar *et al*., 2019) (Supplementary Table S2), which were estimated by Splatter from 8,000 cells of the organoid dataset in Quadrato et al. (Quadrato *et al*., 2017). We simulated two batches and seven cell types. The number of cells in each batch was 2,000, and the number of genes was 10,000. Ten baseline datasets with balanced settings using different seeds were first simulated. Then, we considered three general scenarios, which were unbalanced batch size (scenario 1), rare cell types (scenario 2), and batch-specific cell types (scenario 3). For each scenario, we further considered three sub-levels. For scenario 1, cells in batch 1 were downsampled to 50% (batch1-0.5), 25% (batch1-0.25), and 12.5% (batch1-0.125); for scenario 2, cells labeled as Group1 in each batch were downsampled to 50% (rare1-0.5), 20% (rare1-0.2), and 10% (rare1-0.1); for scenario 3, the number of common cell types were reduced to five (common-5), three (common-3), and one (common-1) separately. Therefore, nine other datasets were generated based on each baseline dataset, and there were in total 10 sub-scenarios, and each of them contained 10 random repeats.

We considered 10 single-cell datasets to investigate the performance of ResPAN in comparisons with other methods, including eight small-to-moderate-scale datasets and two large-scale datasets. In addition, we used a dataset from patients with colorectal cancer to demonstrate the ability of ResPAN on downstream analysis. The sources of these datasets can be found in Supplementary Table S3. The cell type composition of each dataset can be found in Supplementary Table S4.

All datasets were preprocessed using the Scanpy pipeline (Wolf *et al*., 2018). Specifically, cells with less than 200 genes expressed and genes expressed in less than three cells were removed. Then, we performed library size normalization and *log*_+1_ transformation on count data. Finally, the top 2,000 HVGs were selected by controlling for the relationship between dispersion and average expression (Satija *et al*., 2015).

### 2.3 Benckmarking process

We benchmarked ResPAN against seven other methods (Seurat v4, iMAP, Harmony, scVI, MNN, Liger, and BBKNN) that have been shown to have good performance on batch correction in previous studies (Luecken *et al*., 2021; Tran *et al*., 2020). We used UMAP (McInnes *et al*., 2018) as a visualization tool to qualitatively evaluate the results. Moreover, to better assess the performance of different methods on both batch correction and biological information conservation, we considered metrics under the two categories from previous studies. There are five metrics that evaluate batch mixing, including 1 minus batch ASW (1-bASW) (Rousseeuw, 1987), batch LISI (bLISI) (Korsunsky *et al*., 2019), kBET (Büttner *et al*., 2019), graph connectivity (Luecken *et al*., 2021), and true positive rate (TP rate) (Wang *et al*., 2021a). Meanwhile, there are 11 metrics that quantify the conservation of biological information from different aspects, including cell type ASW (cASW), 1 minus cell type LISI (1-cLISI), NMI (Pedregosa *et al*., 2011), ARI (Hubert and Arabie, 1985), positive rate (Wang *et al*., 2021a), cell cycle score (CC score) (Luecken *et al*., 2021), HVG score (Luecken *et al*., 2021), kNN similarity (kNN sim), cell-cell similarity preservation (cell-cell sim), expression similarity (exp sim), and differentially expressed gene detection F1 score (DEG F1). cASW, 1-cLISI, NMI, ARI, and positive rate quantify the preservation of cell clusters before and after batch correction and require true cell type labels as inputs. For CC score, HVG score, kNN sim, cellcell sim pres, exp sim, and DEG F1 score, they are label-free metrics and evaluate the preservation of cell cycle variance, top HVGs, kNNs, cell-cell similarity matrix, and expression levels, respectively. Label-free metrics were calculated for each batch and final scores were averaged across batches. For the calculation of DEG F1 score on simulated data, DEGs detected after correction are compared with true DEGs. All metrics are ranged from 0 to 1, with higher value indicating better performance.

Among all the metrics, only HVG score, exp sim, and DEG F1 score require corrected data in the gene space. For other metrics that require embedded data, we took the top 20 PCs of the integrated data when a method generates corrected gene expression data (ResPAN, iMAP, MNN, and Seurat v4). For methods that only generate corrected data in an embedding space (Liger, scVI, and Harmony), we set the dimension as 20 and took the corrected embedded data to calculate those metrics. For BBKNN, since this method does not generate integrated gene expression data or latent representations, we did not compute metrics for this method. Furthermore, we removed batch-specific cell types when calculating metrics related to batch mixing. To overcome different batch distributions across cell types, all batch-related metrics were calculated in a cell-type-specific manner. For the deep learning based models (ResPAN, iMAP, and scVI), due to the stochasticity in the optimization algorithm, we ran them on each real dataset using 10 different seeds and took the average when calculating the metrics mentioned above. For simulation data, since each scenario contained 10 random datasets, we ran those methods only once for each data using a fixed seed, but took the average across the 10 repeated datasets to evaluate their performance under the corresponding scenario.

After getting each metric, a batch correction score and a bio conservation score were calculated by averaging all metrics under the respective category. Then, we took the weighted average of the batch score and the bio score as a measurement of the overall performance of a model. We used the same weight (0.6 for batch and 0.4 for bio) as used in a newly published benchmarking paper (Luecken *et al*., 2021) to put more emphasis on biological information conservation. Visualizations of quantitative evaluations in this paper were obtained by re-implementing the plotting functions provided by the same benchmarking paper, which was inspired by the code of Saelens et al. (Saelens *et al*., 2019).

Detailed description of all metrics can be found in Supplementary Note S1.3. The versions of software packages we used can be found in Supplementary Table S5. The device information of our experiments can be found in Supplementary Note 1.4.

## 3 Results

### 3.1 Simulation study

From UMAP visualizations in Fig. 2a and Supplementary Fig. S1, we can see that ResPAN was able to remove the batch effect in all the 10 scenarios while maintaining the separation of different simulated cell clusters. In terms of quantitative evaluations (Fig. 2b), ResPAN achieved the highest overall score by taking an average across all 10 scenarios, demonstrating its ability to maintain a good balance between batch mixing and bio conservation across different scenarios. Harmony performed worst in the simulation study with low scores on both batch correction and bio conservation. Although Liger and scVI were the top methods in terms of batch correction score, they performed poorly on bio conservation. On the contrary, MNN and Seurat v4 achieved higher bio conservation scores but lower batch correction scores. As for iMAP, its performance was moderate in terms of both batch correction and bio conservation, and only ranked fifth among all methods. The breakdown of all metrics in each of the 10 scenarios can be found in Supplementary Table S6.

**Figure 2:**
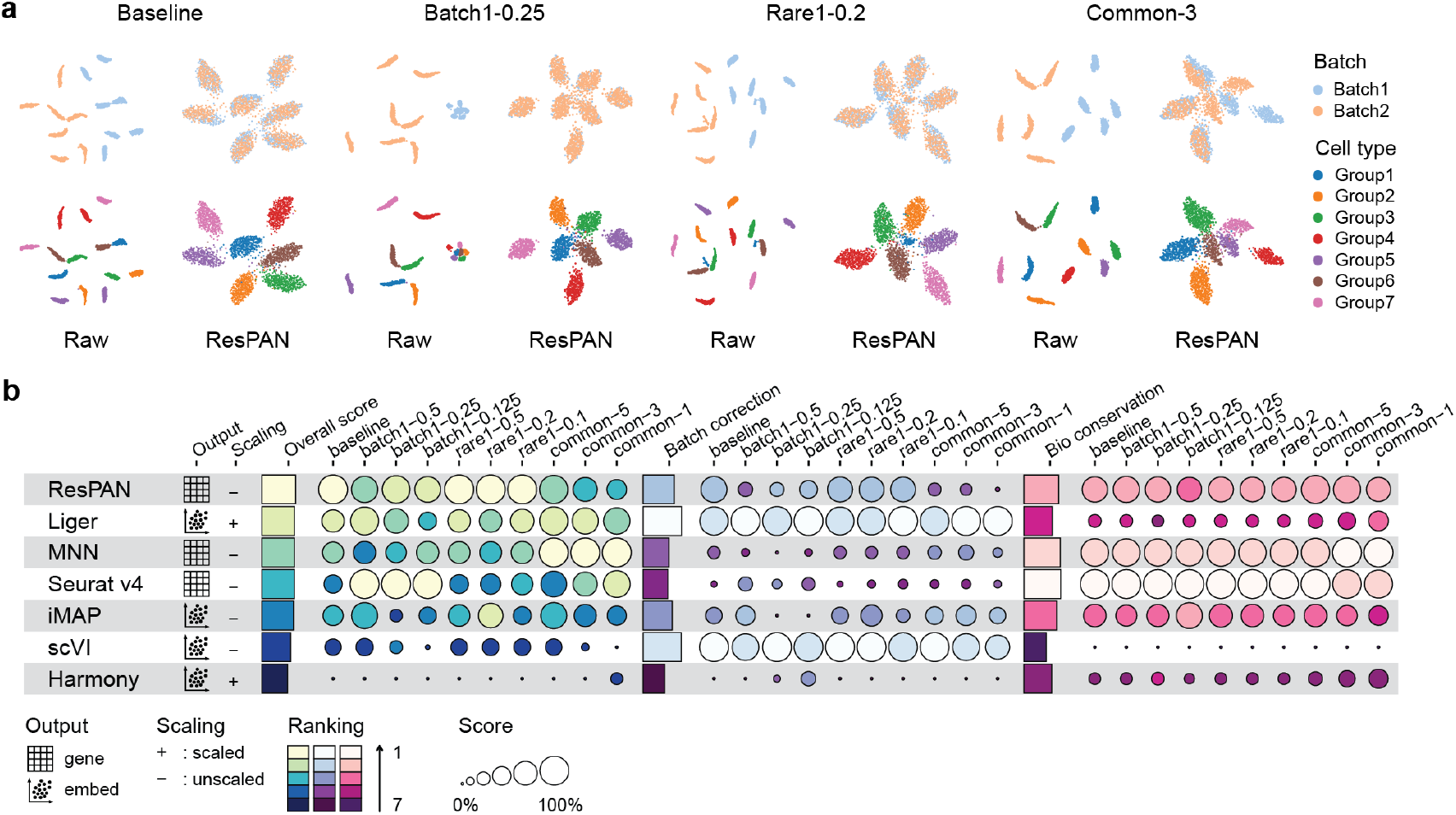
Benchmarking results on simulation data. (a) UMAP visualization of raw data and ResPAN under baseline, batch1-0.25, rare1-0.2, and common-3. For other scenarios and other methods, please see Supplementary Fig. S1 (b) An overview of the quantitative evaluation. Metrics are divided into three parts, bio conservation, batch correction, and overall score (a 40/60 weighted mean of batch and bio score under each scenario). For each part, detailed information of each scenario is shown in circles, overall information is represented by bars after taking an average over all scenarios. Methods are ranked by overall score.

Apart from comparing ResPAN’s performance against other batch correction methods, we also investigated whether changing full skip connection on the generator model to one skip connection on the output layer only (this is the generator model used in iMAP) could affect the performance. From Supplementary Fig. S2a, we can observe that the design of a fully skip-connected generator achieved higher batch correction scores, while the only exception was the common-1 scenario, where only one out of seven cell types is shared among batches. Such trend did not exist in the bio conservation scores, but the overall scores showed similar patterns as the batch scores.

### 3.2 Benchmark on real scRNA-seq data

For real data benchmarking, we first compared our model against other methods on datasets with known cell type labels and small-to-moderate data sizes (less than 100,000 cells). The eight datasets we chose are pure cell lines (CL) (Zheng *et al*., 2017), human pancreas (Wang *et al*., 2021a), human peripheral blood mononuclear cells (Human PBMC) (Zheng *et al*., 2017), PBMC 3&68K (Zheng *et al*., 2017), mouse hematopoietic stem and progenitor cells (MHSP) (Nestorowa *et al*., 2016; Paul *et al*., 2015), Mouse Cell Atlas (MCA) (Consortium *et al*., 2018; Han *et al*., 2018), Mouse Retina (Macosko *et al*., 2015; Shekhar *et al*., 2016), and a multi-batch human lung dataset (Vieira Braga *et al*., 2019). In the following paragraph, we will use the CL dataset as an example to compare ResPAN against other methods.

From Fig. 3a, we can observe that ResPAN delivered the most satisfactory visualization. Among other methods, MNN and Seurat V4 failed to integrate 293T cells, and Liger and BBKNN’s integration results of Jurkat cells were not complete. Moreover, for scVI, there seems to exist mixing of Jurkat and 293T cells around the boundary, and Harmony did not generate homogeneous integrated Jurkat cells. From the metrics (Fig. 3b and 3c), ResPAN performed very well on both batch correction and bio conservation, and achieved the highest overall score (24.25% higher than the second-best-performer Harmony). One thing worth mentioning is that although iMAP delivered a good visualization in Fig. 3a, its scores were the worst in Fig. 3c. This is because we ran all neural network methods, including iMAP, 10 times using different seeds. In addition, iMAP’s performance varied a lot under different seeds, indicating the unstability of iMAP. UMAP visualizations under different seeds for ResPAN, iMAP and scVI are shown in Supplementary Fig. S3, and iMAP totally failed in six out of the 10 cases. This observation was not only found in the CL dataset (Supplementary Fig. S4). For other datasets, the UMAP visualizations and two-coordinate representations of batch score and bio score can be found in Supplementary Fig. S5 and S6, respectively. We did not detect any obvious flaws of ResPAN in integrating these datasets, and ResPAN’s dot was located in the upper right part of the two-coordinate representations of most datasets.

**Figure 3:**
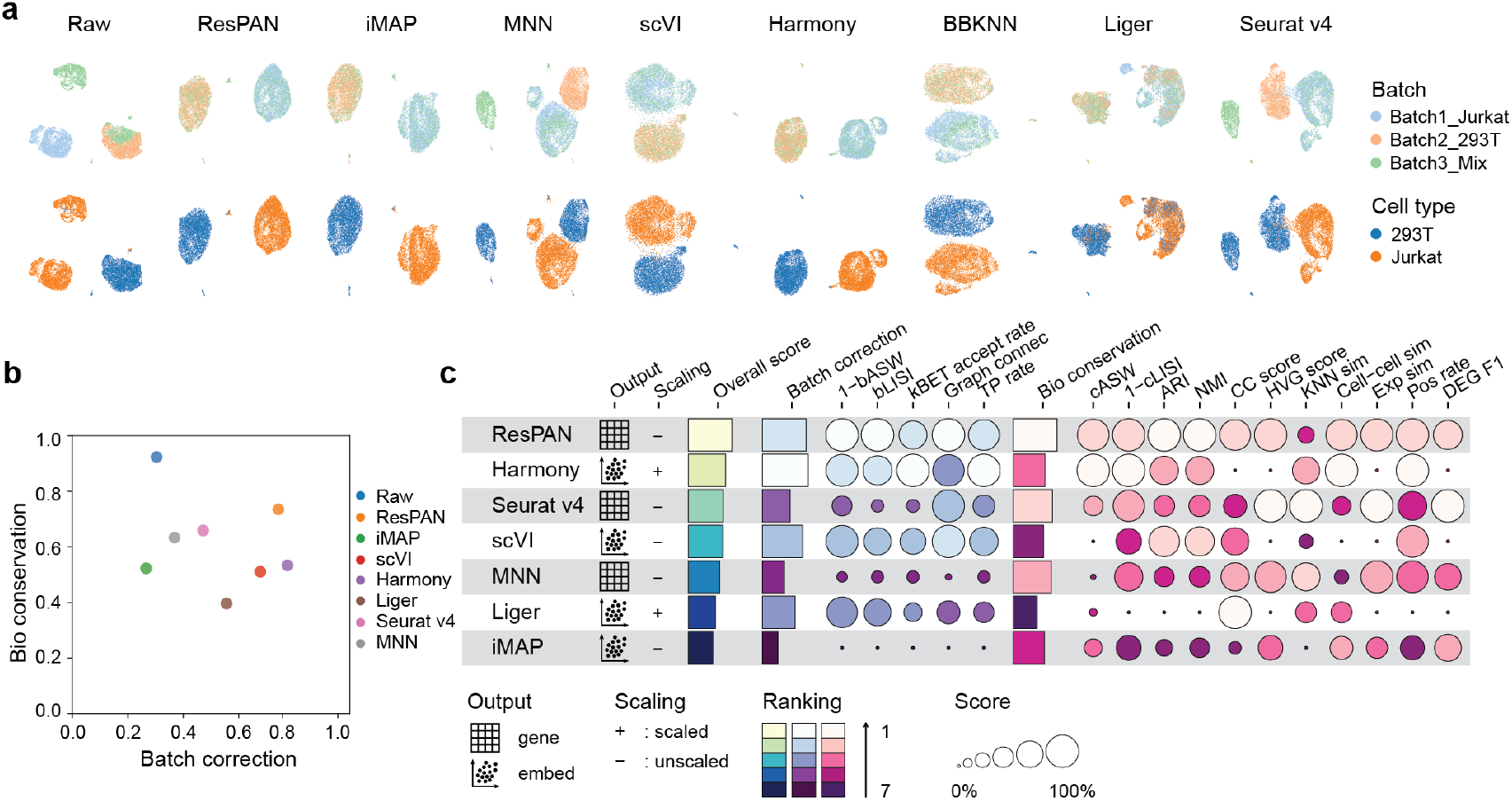
Benchmarking results on the CL dataset. (a) UMAP Visualizations of raw data and eight methods. (b) Bio conservation score versus batch correction score. Both scores range between 0 and 1, with larger value indicating better performance. (c) Detailed breakdown of each metric. Batch correction is calculated as the average of five batch metrics and bio conservation is the average of 11 bio-related metrics. Methods are ranked by overall score, which is a 40/60 weighted mean of batch and bio score.

We recorded the running time and peak memory usage of different methods in the eight datasets (Supplementary Table S7, sheet named ‘Time’ and ‘Memory’, datasets placed in descending order of their cell numbers). We found that MNN was the most time-consuming method, especially for the two medium-scale datasets, Lung (about 30,000 cells) and Mouse Retina (about 70,000 cells). It took MNN about 16 hours and two hours to run on them, respectively. Moreover, MNN required more memory than all the other methods on these two datasets. BBKNN and Harmony were the two fastest methods for running on both small-scale and medium-scale datasets. This might be because their implementations do not involve large matrix decomposition as Liger or nearest neighbor construction on highdimensional space as MNN does. For Liger and Seurat v4, although they were able to finish in several minutes on small-scale datasets, their running time increased to about an hour on medium-scale datasets. For small-scale datasets, we found that BBKNN, Harmony, Liger, and Seurat v4 consistently took less memory than the three neural network methods (ResPAN, iMAP and scVI). As for the neural network methods, their running time and memory usage remained almost constant as data size increased and were among the fastest and most memory-efficient methods on medium-scale datasets. They were able to finish in less than 10 minutes, except for ResPAN and iMAP on Lung. ResPAN and iMAP ran longer than scVI on Lung because they were based on sequential training by mapping each batch successively to the reference data, and Lung contains the most number of batches (16). However, the running time was still less than half an hour.

The scalability of a computational method has become more important, as singlecell studies usually sequence hundreds of thousands or even millions of cells and the analysis of large-scale datasets can take much space, memory, and time, making batch effect removal even more challenging. Therefore, we further considered two large-scale scRNA-seq datasets with more than 500,000 cells and investigated the ability of ResPAN to integrate large-scale datasets. The first dataset is a Mouse Brain dataset (Rosenberg *et al*., 2018; Saunders *et al*., 2018), which contains over 800,000 cells. The second dataset is from the Human Cell Atlas (HCA) (Regev *et al*., 2017) and contains more than 700,000 cells.

As can be seen from Fig. 4a and 4b, although these two large-scale datasets have significant batch effects, ResPAN successfully removed the batch effects on both of them, and its performance was as good as another best-performing method, Seurat v4. ResPAN also achieved a good balance between batch correction and bio conservation on these two datasets (Fig. 4c). Apart from fulfilling the batch correction task, the running time and memory usage of an algorithm are another set of important indicators of its efficiency and scalability on large-scale datasets. As shown in Table 1, ResPAN along with iMAP and scVI were the fastest algorithms on the two large-scale datasets and their memory usage was as efficient as two traditional methods, Harmony and BBKNN. This is because all these three methods are based on neural networks, and the optimization of neural networks is performed through mini-batch training and stochastic gradient descent (SGD). Among them, ResPAN was the fastest one and took less memory than iMAP, which was also observed in small-to-medium-scale datasets (Supplementary Table S7). Compared to iMAP, which performs MNN pairing on a latent space learned by training an additional complicated autoencoder, ResPAN has a lighter structure and searches MNN pairs through PCA-reduced space. This helps reduce running time and memory usage without sacrificing its good performance on both batch correction and bio conservation. Another thing worth mentioning is that although Seurat v4 performed as well as ResPAN in terms of UMAP visualizations and quantitative metrics, it took around three days and more than 800 GB of memory to finish. Thus, ResPAN should be the most recommended method for large-scale datasets in real practice.

**Figure 4:**
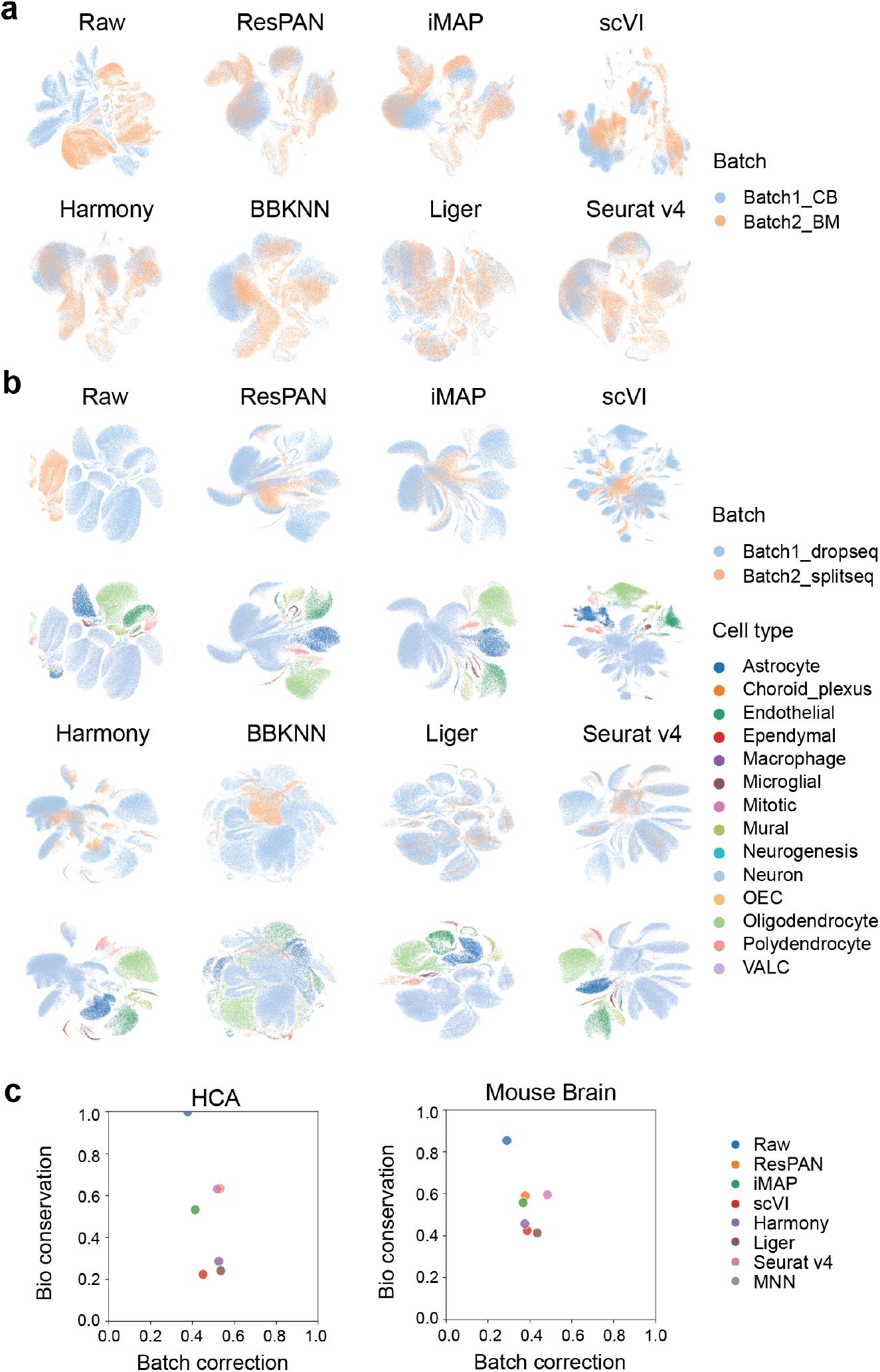
Benchmarking results on large-scale datasets. (a) UMAP visualizations of the HCA dataset across different methods. (b) UMAP visualizations of the Mouse Brain dataset across different methods. (c) Bio conservation score versus batch correction score for the HCA (left) and the Mouse Brain (right) dataset. OEC: olfactory ensheathing cells; VALC: vascular and leptomeningeal cells.

**Table 1:** Running time and peak memory usage on large-scale datasets

| Dataset | ResPAN | iMAP | MNN | scVI | Harmony | Liger | Seurat v4 | BBKNN |
| --- | --- | --- | --- | --- | --- | --- | --- | --- |
| **Running time** |  |  |  |  |  |  |  |  |
| Mouse Brain (unit: h) | 0.0491 | 0.0649 | TIMEOUT | 0.158 | 1.49 | 8.96 | 74.7 | 0.111 |
| HCA (unit: h) | 0.0487 | 0.0589 | TIMEOUT | 0.147 | 0.643 | 44.8 | 68.9 | 0.0824 |
| **Peak memory Usage** |  |  |  |  |  |  |  |  |
| Mouse Brain (unit: GB) | 35.89 | 42.77 | TIMEOUT | 28.78 | 26.56 | 251.07 | 853.36 | 26.99 |
| HCA (unit: GB) | 29.18 | 34.69 | TIMEOUT | 20.23 | 19.65 | 209.82 | 886.44 | 19.89 |

At last, we combined all metrics calculated on both the eight small-to-moderate-scale datasets and the two large-scale datasets to have an overall assessment of the performance on real data. As shown in Fig. 5, ResPAN achieved the highest overall score by taking all 10 datasets into consideration, and it was always among the top-three methods across all datasets. It was the top-ranked model in four datasets (CL, PBMC 3&68k, MCA, HCA), and was the second-ranked model in five datasets (Pancreas, Human PBMC, MHSP, Mouse Retina, Mouse Brain). Detailed breakdown of each metric in all real datasets can be found in Supplementary Table S7.

**Figure 5:**
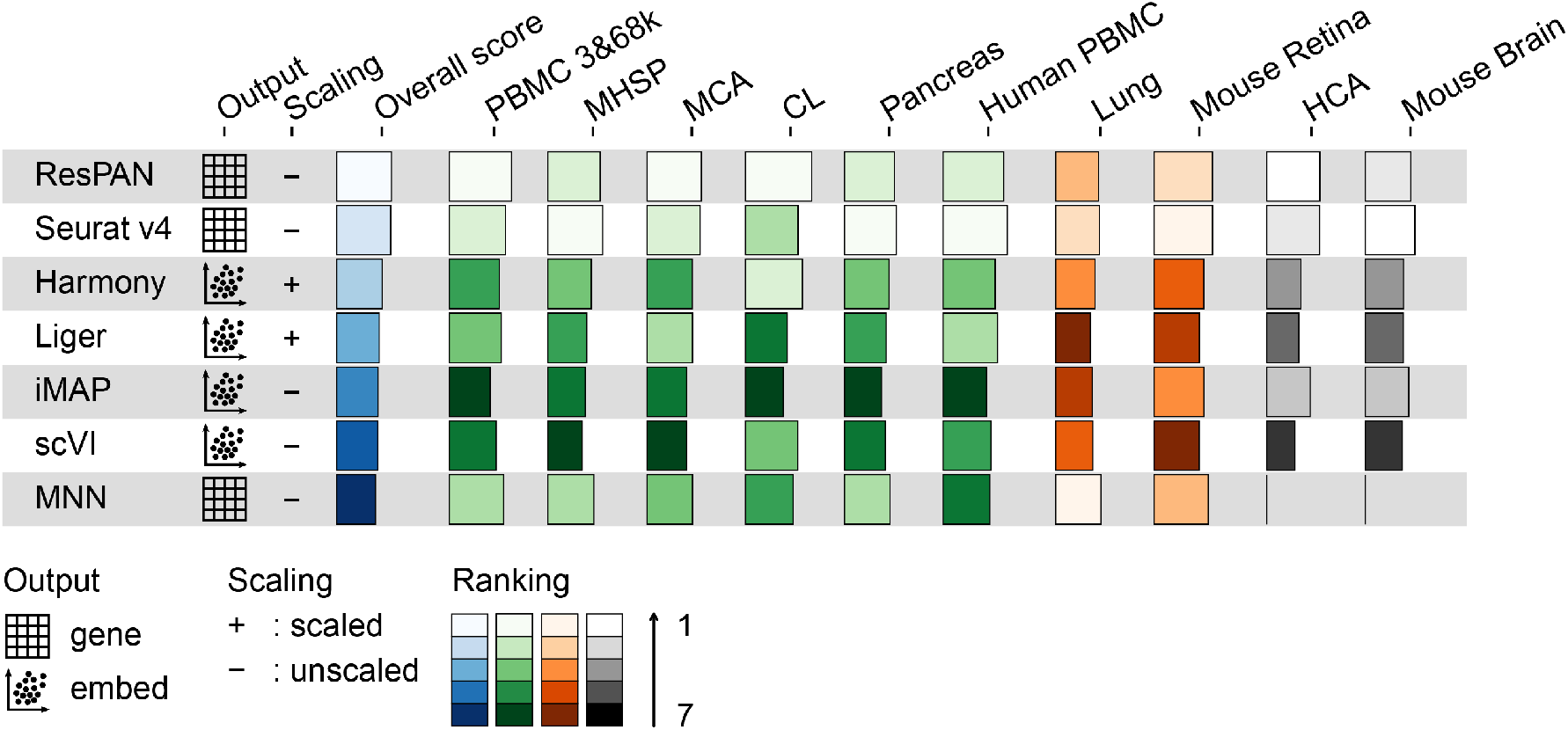
Overview of quantitative evaluations on real data. Green bars, orange bars, and grey bars are the overall scores for the six small-scale, the two medium-scale, and the two large-scale datasets across different methods, respectively. The ‘Overall score’ in blue bars are calculated by taking the mean of overall scores across 10 datasets. Datasets are arranged in the increasing order of their size from left to right.

As we did in the simulation study, we compared the performance of ResPAN using a full skip connection versus using one skip connection on the output layer of the generator on real data. From Supplementary Fig. S2b, we can observe a similar trend as observed in the simulation study, which is that the design of a fully skip-connected generator achieved higher batch correction scores consistently across all data.

### 3.3 Application to a Colorectal Cancer data

The Colorectal Cancer (CRC) dataset is a scRNA-seq dataset containing 54,285 tumor-infiltrating immune cells from 18 CRC patients using both Smart-seq2 and 10x platforms (Zhang *et al*., 2020). Here, we take a deeper dive into the functional diversity of ResPAN by looking into the CRC dataset. Results shown in Fig. 6a suggest that ResPAN could mostly eliminate batch effects in the CRC dataset and preserve the biological variation of the dataset. From Fig. 6a, we note that innate lymphoid cells (ILCs) and CD8 T cells were close to each other in the UMAP after removing batch effects. We assumed that ILCs were mixed with CD8 T cells. To verify this assumption, based on the traceplot clustered by cell type labels (Fig. 6b), we found that the gene expression levels between CD8 T cells and ILCs were very similar. One possible reason is that some ILCs might be incorrectly marked. This could be caused by the expression levels of some maker genes being too close or noises in the original data. Another possible reason is that the two types of cells could convert to each other.

**Figure 6:**
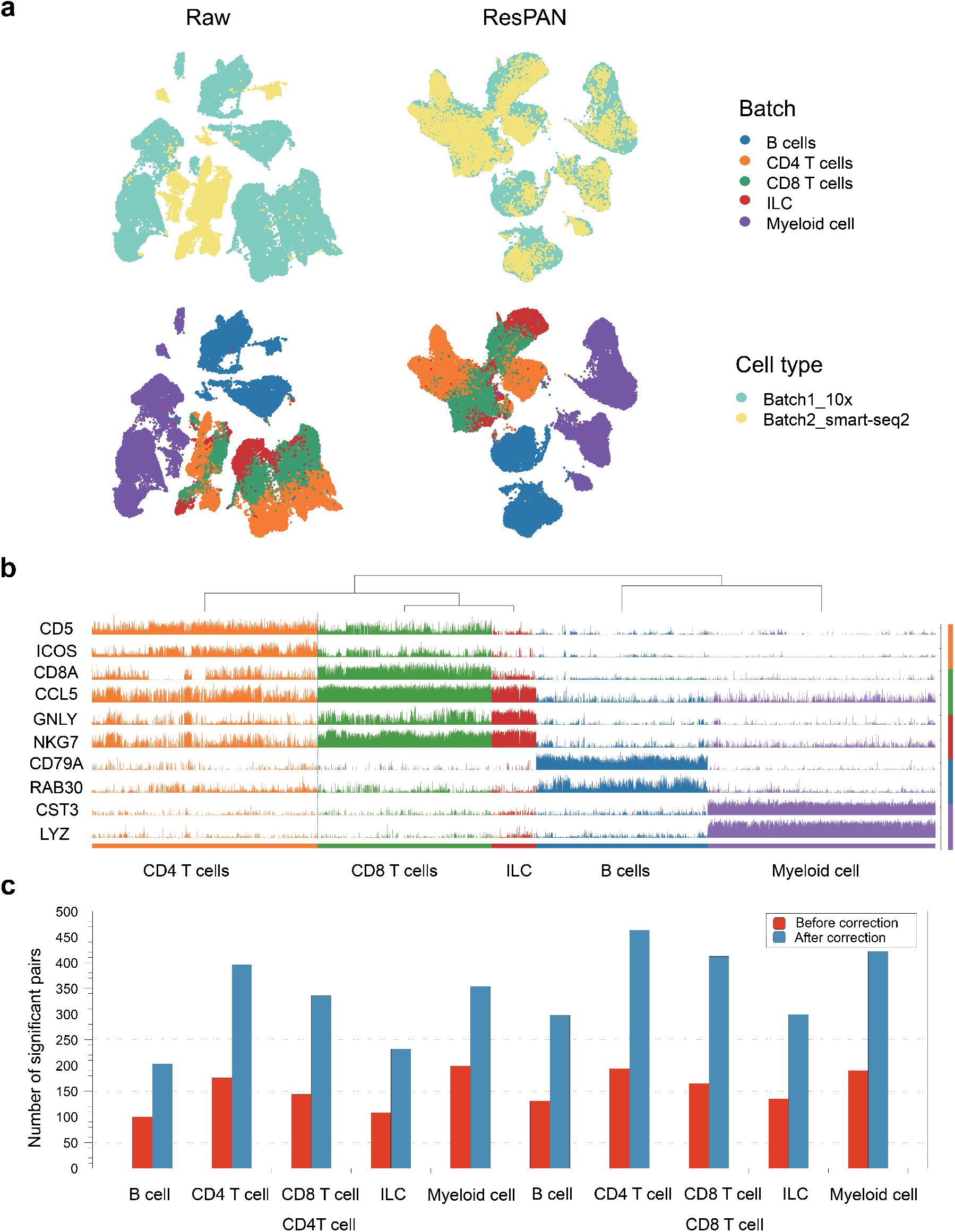
Application of ReaPAN on the CRC dataset. (a) UMAP visualizations before (first column) and after batch correction by ResPAN (second column). (b) Normalized gene expression traceplot of the CRC dataset after batch effect removal. (c) Number of significant ligand-receptor pairs (*p* < 0.05) defined between all other cell types and T cells.

We also studied the interactions among different cell types. The two batches of the CRC dataset were sequenced based on different platforms, and the two platforms had their own characteristics. The 10x data (10x original) are sparser but have higher throughput in terms of cell numbers, while the Smart-seq2 data contain more accurate measurement of gene expression levels (Wang *et al*., 2021a,b; Zhang *et al*., 2020). Therefore, relying on the noisy 10X data only may make it difficult to estimate the correlation between cells and calculate cell-cell communication scores. ResPAN provides a potential solution to this problem by mapping the 10x data to the Smart-seq2 data. We used CellPhoneDB (Efremova *et al*., 2020) to calculate cell-cell communication scores and generate visualization results. Based on Fig. 6c, we found that more ligand-receptor pairs were identified using the 10x data after batch correction (10x correct), which could be defined as potential novel cellcell interactions. To verify that our selected pairs are meaningful, we obtained the ligand-receptor pairs from the Smart-seq2 dataset as the ‘ground truth’ and calculated the overlap score (range: [0,1]) between this dataset and 10x correct dataset. The score was as high as 0.87, which was higher than the score of iMAP (0.76) and Seurat v4 (0.80). Therefore, the corrected data generated by ResPAN were the closest one to the Smart-seq2, and ResPAN has the potential to uncover novel cell-cell interactions. In Supplementary Fig. S7, we show some of the newly discovered pairs. The number of significant ligand-receptor pairs generated based on iMAP and Seurat v4 can be found in Supplementary Fig. S8.

## 4 Discussion

In this paper, we have proposed ResPAN, a light-structured residual adversarial network model guided by MNN pairs for removing batch effects. We designed a comprehensive benchmarking pipeline for evaluating ResPAN and seven other competitors through both simulation study and real data assessment. In total, we considered 16 metrics, among which five quantify the batch mixing and 11 evaluate the preservation of biological information. Among the 11 bio-related metrics, only five are traditional metrics that require cell labels and were used extensively in previous studies (Tran *et al*., 2020; Wang *et al*., 2021a) for benchmarking different batch correction models. The remaining six metrics are all label free and consider biological information from additional aspects, such as cell cycle variations, HVGs, DEGs, and cell-cell relationships. We argue that the current framework is able to reflect the performance of a batch correction method more comprehensively and conclusions drawn from all these scores will not be biased towards any specific ones.

Based on the overall score calculated by first taking a weighted mean of bio and batch scores followed by averaging across all scenarios (simulation) or datasets (real), we found that ResPAN was the top performer on both simulated and real data. The other two top-performers on simulated data were Liger and MNN, demonstrating the suitable performance of these two methods on less heterogeneous data but more unbalanced data settings. However, the other two top-performers on 10 real datasets were totally different, which were Seurat v4 and Harmony. Thus, ResPAN was the only method that could generate desirable results for all types of data. This should thank to the nature of generative adversarial training and the MNN-based training data construction. With a framework based on GANs, ResPAN is able to map the query data onto the reference data without explicitly specifying any distribution family or a single batch variable as scVI does. This ensures the flexibility and the generalization ability of ResPAN across various datasets and scenarios. The MNN based data generation further guarantees that batch-specific biological information can be maintained since query cells distributed largely differently from reference cells would not be selected for training and vice versa. In brief, GAN works as a powerful tool for finding the correct mapping that removes the batch effect, and MNN-based data generation works as a gatekeeper to prevent batch-specific biological information from being removed by the adversarial network.

Although iMAP is also a GAN based batch correction model, its average performance was much worse than ResPAN on both simulated and real data. This is because iMAP totally failed to remove batch effects under some random seeds (Supplementary Fig. S3 and S4). There are two major differences in the design of ResPAN and iMAP. First, iMAP performs MNN pairing on a latent space learned by training an additional complicated autoencoder, while ResPAN has a lighter structure and searches MNN pairs through PCA-reduced space. Second, we use a fully skip-connected adversarial autoencoder as the generator, while iMAP only uses skip connection for the output layer. The second design was found to consistently improve batch correction scores on all real data and the majority of simulated data (Supplementary Fig. S2). However, removing such structures did not cause failures as experienced by iMAP. Therefore, the first difference might be the reason for the unstability of iMAP, but we can not rule out the possibility of coding issues in iMAP. Another drawback of iMAP’s cumbersome design is that it takes more time and memory for iMAP to run than ResPAN, especially when dealing with large-scale datasets.

The average overall score of Seurat v4 (0.598) was very close to ResPAN’s (0.608) on real datasets. However, we observed failure in the batch correction of the CL dataset for Seurat v4. In contrast, we did not observe any failures on real datasets for ResPAN. Moreover, when running on large datasets, ResPAN was able to finish within five minutes and required memory of 35 GB, while Seurat v4 ran for almost three days with peak memory usage of over 800 GB. This is super inefficient and inconvenient for researchers to apply it to large-scale data. Moreover, for Harmony, scVI, Liger, and BBKNN, they do not return corrected gene expression data. Instead, Harmony, scVI, and Liger return corrected data in a low dimensional space and BBKNN only modifies the construction of the nearest neighborhood graph. In contrast, methods like ResPAN, MNN, Seurat v4 and iMAP can generate corrected gene expression data and provide a better basis for downstream analysis. For example, in the CRC dataset, ResPAN showed its ability to identify potentially misclassified cells and uncover hidden cell-cell interactions.

## 5 Conclusion

In summary, we demonstrate that ResPAN is a powerful method for batch effect removal through benchmarking studies on both simulated and real data across different scenarios and scales in comparison with seven other batch effect removal methods using 16 metrics. Firstly, ResPAN can achieve a good balance between batch correction and biological information conservation. It was the top performer based on the overall score on both simulated and real data. Secondly, ResPAN is a flexible model and researchers can adapt many hyper-parameters in our model to their needs. Meanwhile, the default parameters could deliver good results, as used in our study. The strong adaptability of our model also allows it to be extended and improved to achieve better performance. Thirdly, ResPAN has a lighter structure and takes the advantages of deep learning models through mini-batch training and SGD, which make it scalable to large-scale datasets while maintaining good performance. We believe that our model can be a powerful and efficient tool for data integration and can aid in better informing downstream analyses, such as cell-type specific biological function analysis, cell-cell heterogeneity analysis, and others.

## References

Arjovsky, M. et al. (2017). Wasserstein generative adversarial networks. In International conference on machine learning, pages 214–223. PMLR.

Büttner, M. et al. (2019). A test metric for assessing single-cell rna-seq batch correction. Nature methods, 16(1), 43–49.

Consortium, T. M. et al. (2018). Single-cell transcriptomics of 20 mouse organs creates a tabula muris. Nature, 562(7727), 367–372.

Efremova, M. et al. (2020). Cellphonedb: inferring cell–cell communication from combined expression of multi-subunit ligand–receptor complexes. Nature protocols, 15(4), 1484–1506.

Gulrajani, I. et al. (2017). Improved training of wasserstein gans. In I. Guyon, U. V. Luxburg, S. Bengio, H. Wallach, R. Fergus, S. Vishwanathan, and R. Garnett, editors, Advances in Neural Information Processing Systems, volume 30. Curran Associates, Inc.

Haghverdi, L. et al. (2018). Batch effects in single-cell rna-sequencing data are corrected by matching mutual nearest neighbors. Nature biotechnology, 36(5), 421–427.

Han, X. et al. (2018). Mapping the mouse cell atlas by microwell-seq. Cell, 172(5), 1091–1107.

Heusel, M. et al. (2017). Gans trained by a two time-scale update rule converge to a local nash equilibrium. Advances in neural information processing systems, 30.

Hornik, K. et al. (1989). Multilayer feedforward networks are universal approximators. Neural networks, 2(5), 359–366.

Hubert, L. and Arabie, P. (1985). Comparing partitions. Journal of classification, 2(1), 193–218.

Johnson, W. E. et al. (2007). Adjusting batch effects in microarray expression data using empirical bayes methods. Biostatistics, 8(1), 118–127.

Korsunsky, I. et al. (2019). Fast, sensitive and accurate integration of single-cell data with harmony. Nature methods, 16(12), 1289–1296.

Kotliar, D. et al. (2019). Identifying gene expression programs of cell-type identity and cellular activity with single-cell rna-seq. Elife, 8, e43803.

Leek, J. T. et al. (2010). Tackling the widespread and critical impact of batch effects in high-throughput data. Nature Reviews Genetics, 11(10), 733–739.

Lopez, R. et al. (2018). Deep generative modeling for single-cell transcriptomics. Nature methods, 15(12), 1053–1058.

Luecken, M. D. et al. (2021). Benchmarking atlas-level data integration in single-cell genomics. Nature Methods, pages 1–10.

Macosko, E. Z. et al. (2015). Highly parallel genome-wide expression profiling of individual cells using nanoliter droplets. Cell, 161(5), 1202–1214.

McInnes, L. et al. (2018). Umap: Uniform manifold approximation and projection for dimension reduction. arXiv preprint arXiv:1802.03426.

Nestorowa, S. et al. (2016). A single-cell resolution map of mouse hematopoietic stem and progenitor cell differentiation. Blood, The Journal of the American Society of Hematology, 128(8), e20–e31.

Papalexi, E. and Satija, R. (2018). Single-cell rna sequencing to explore immune cell heterogeneity. Nature Reviews Immunology, 18(1), 35.

Paul, F. et al. (2015). Transcriptional heterogeneity and lineage commitment in myeloid progenitors. Cell, 163(7), 1663–1677.

Pedregosa, F. et al. (2011). Scikit-learn: Machine learning in python. the Journal of machine Learning research, 12, 2825–2830.

Pijuan-Sala, B. et al. (2018). Single-cell transcriptional profiling: a window into embryonic cell-type specification. Nature Reviews Molecular Cell Biology, 19(6), 399–412.

Polański, K. et al. (2020). Bbknn: fast batch alignment of single cell transcriptomes. Bioinformatics, 36(3), 964–965.

Quadrato, G. et al. (2017). Cell diversity and network dynamics in photosensitive human brain organoids. Nature, 545(7652), 48–53.

Regev, A. et al. (2017). Science forum: the human cell atlas. elife, 6, e27041.

Ritchie, M. E. et al. (2015). limma powers differential expression analyses for rna-sequencing and microarray studies. Nucleic acids research, 43(7), e47–e47.

Rosenberg, A. B. et al. (2018). Single-cell profiling of the developing mouse brain and spinal cord with split-pool barcoding. Science, 360(6385), 176–182.

Rousseeuw, P. J. (1987). Silhouettes: a graphical aid to the interpretation and validation of cluster analysis. Journal of computational and applied mathematics, 20, 53–65.

Saelens, W. et al. (2019). A comparison of single-cell trajectory inference methods. Nature biotechnology, 37(5), 547–554.

Satija, R. et al. (2015). Spatial reconstruction of single-cell gene expression data. Nature biotechnology, 33(5), 495–502.

Saunders, A. et al. (2018). Molecular diversity and specializations among the cells of the adult mouse brain. Cell, 174(4), 1015–1030.

Shaham, U. et al. (2017). Removal of batch effects using distribution-matching residual networks. Bioinformatics, 33(16), 2539–2546.

Shekhar, K. et al. (2016). Comprehensive classification of retinal bipolar neurons by single-cell transcriptomics. Cell, 166(5), 1308–1323.

Stuart, T. et al. (2019). Comprehensive integration of single-cell data. Cell, 177(7), 1888–1902.

Suvà, M. L. and Tirosh, I. (2019). Single-cell rna sequencing in cancer: lessons learned and emerging challenges. Molecular cell, 75(1), 7–12.

Tran, H. T. N. et al. (2020). A benchmark of batch-effect correction methods for single-cell rna sequencing data. Genome biology, 21(1), 1–32.

Vieira Braga, F. A. et al. (2019). A cellular census of human lungs identifies novel cell states in health and in asthma. Nature medicine, 25(7), 1153–1163.

Wang, D. et al. (2021a). imap: integration of multiple single-cell datasets by adversarial paired transfer networks. Genome biology, 22(1), 1–24.

Wang, X. et al. (2021b). Direct comparative analyses of 10x genomics chromium and smart-seq2. Genomics, Proteomics & Bioinformatics.

Welch, J. D. et al. (2019). Single-cell multi-omic integration compares and contrasts features of brain cell identity. Cell, 177(7), 1873–1887.

Wolf, F. A. et al. (2018). Scanpy: large-scale single-cell gene expression data analysis. Genome biology, 19(1), 1–5.

Zappia, L. et al. (2017). Splatter: simulation of single-cell rna sequencing data. Genome biology, 18(1), 1–15.

Zhang, L. et al. (2020). Single-cell analyses inform mechanisms of myeloid-targeted therapies in colon cancer. Cell, 181(2), 442–459.

Zheng, G. X. et al. (2017). Massively parallel digital transcriptional profiling of single cells. Nature communications, 8(1), 1–12.

