## Supplementary Notes for "ResPAN: a powerful batch correction model for scRNA-seq data through residual adversarial networks"

### 1 Methods

#### 1.1 Random walk MNN based training data generation

As we discussed in the main manuscript, Random walk MNN (rwMNN) pairs are generated on a latent space (top 20 principal components) to get the training data for ResPAN. Here, we describe the details of this process below.

We observe that most studies on batch effect removal chose a gene number within the range of at least 200 and up to 2000 (Tran *et al.*, 2020; Wang *et al.*, 2021) based on the highly variable gene (HVG) filtering method (Satija *et al.*, 2015). In this paper, we follow the same rule to process the raw count matrix. After obtaining the filtered count matrix, we proceed to the step of generating the training set. Firstly, we utilize principal component analysis (PCA) to generate latent space representation of gene expression data to perform data denoising. Inspired by the basic attention mechanism, we construct mutual nearest neighbor (MNN) cell pairs (Haghverdi *et al.*, 2018) using cosine similarity based on the latent space data. We generate larger training dataset by using random-walk based oversampling strategy. That is, we use MNN pairs between reference dataset and query dataset as seeds. At the beginning of iteration, k nearest neighbor (kNN) graphs are constructed around cells selected in MNN pairs in the reference data and the query data, and random walk sampling is carried out among kNNs. The sampled pairs are used as seeds for the next round of random walks. This algorithm could record the pairs experienced and add them into the training dataset until enough number of random walks is reached. The number is set as 50 for each MNN pair, so the final number of pairs after finishing random walks should be 50 times that of the number of MNN pairs. iMAP uses a similar scheme and names the pairs as random Walk MNN (rwMNN) pairs, so we continue using that name. Finally, we use the rwMNN pairs as index to obtain the corresponding data on the gene expression space, thus completing the construction of the training data. This iterative algorithm can be found in Algorithm 1. Note that before running the rwMNN algorithm, 3,000 cells are randomly sampled from the reference data and the query data separately if any of them contain more than 3,000 cells. Although as you may observe in Algorithm 1, $k_1$ and $k_2$ are adaptively set based on the data size, we fixed $k_1 = 10$ and $k_2 = 20$ in our study for all simulated and real data.

#### 1.2 The structure and the training of ResPAN

ResPAN is composed of a generator (G) and a discriminator (D). We design our generator in the form of an autoencoder in order to better extract biological information and remove noise from the data. The generator consists of an input layer, five hidden layers and an

**Algorithm 1** rwMNN generating algorithm.

**Input:** Latent representation of reference dataset $LatData_{ref}$; Latent representation of query dataset $LatData_{que}$; Reference dataset $Data_{ref}$; Query dataset $Data_{que}$;
**Output:** Training dataset $Data_{true}$, $Data_{fake}$

INIT: initialize all parameters.
$k_2$ = int(min(len($Data_{ref}$), len($Data_{que}$))/100)
$k_1$ = max(int($k_2$/2), 1)
anchor_pairs = acquire_mnn_pairs($LatData_{ref}$,$LatData_{ref}$, $k_2$)
query_pairs = acquire_mnn_pairs($LatData_{que}$,$LatData_{que}$, $k_2$)
mutual_pairs = acquire_mnn_pairs($LatData_{ref}$,$LatData_{que}$, $k_1$)
anchor_pairs_dict = generate_dict(anchor_pairs)
query_airs_dict = generate_dict(query_pairs)
total_pairs = []
**for** $x, y$ in mutual_pairs **do**
    SeedPoint = $(x, y)$
    **for** *step* in range(0,50) **do**
        total_pairs.append(SeedPoint)
        SeedPoint[0] = RandomChoice(anchor_pairs_dict[SeedPoint[0]])
        SeedPoint[1] = RandomChoice(query_pairs_dict[SeedPoint[1]])
$Data_{true}$ = item[0] for all item in total_pairs
$Data_{fake}$ = item[1] for all item in total_pairs
Return $Data_{true}, Data_{fake}$

output layer. The dimension of the input and the output layer is the number of HVGs. The dimensions of the five hidden layers are 1,024, 512, 256, 512 and 1,024, respectively. The discriminator contains six layers with decreasing dimensions (number of HVGs, 1,024, 512, 256, 128 and 1). For the generator, we also deploy the batch normalization (BN) technique (Ioffe and Szegedy, 2015) to accelerate the convergence speed. According to WGAN-GP (Gulrajani *et al.*, 2017), we do not use BN for the discriminator because we intend to make sure that the smoothness required by the Lipschitz condition is not affected by the BN layers. The loss function of ResPAN is:

$$L = E_{x^* \sim P_g}[D(x^*)] - E_{x' \sim P_r}[D(x')] + \lambda E_{x \sim P_x}[(\|\nabla_x D(x)\|_2 - K)^2], \tag{1}$$

where $P_g$ is the distribution of the data generated from the generator, $P_r$ is the distribution of the reference data, and $P_x$ is the distribution of the data interpolated using the generated ($Fake$) and the reference ($True$) data.

The first two terms represent the Wasserstein distance, which is the original expression in WGAN that we intend to minimize. However, if we minimize this value directly, the K-Lipschitz condition may not be satisfied. The original WGAN enforces the Lipschitz condition by gradient clipping. However, there are still cases where the parameters spontaneously concentrate on the boundary, causing gradient vanishing. WGAN-GP introduces the gradient penalty to solve this problem (Gulrajani *et al.*, 2017). The third term, gradient penalty, first interpolates between true data and fake data to obtain a new distribution. Then, it puts a constraint on the gradient norm of the discriminator to enforce Lipschitz continuity. In our implementation, we take $K = 1$ because as proved by the WGAN-GP paper, the L2 norm of the ideal gradient for WGAN is 1 (1-Lipschitz condition) (Gulrajani *et al.*, 2017). The interpolation function between the $True$ and $Fake$ data is defined as:

$$
\begin{cases} Interpolation(True, Fake) = \epsilon \cdot True + (1-\epsilon) \cdot Fake \\ \epsilon \sim Uniform(0,1) \\ x \sim P_x(Interpolation(True, Fake)) \end{cases} \tag{2}
$$

For the choice of the activation function, we utilize the Mish activation function (Misra, 2019), which is defined as :

$$
Mish(x) = x \cdot tanh(ln(1 + e^x)). \tag{3}
$$

The Mish function has smoother gradients compared with the Rectified Linear Unit (ReLU) (Glorot *et al.*, 2011) function, thus allows better information integration and information mining performed by deep neural networks (Misra, 2019).

For the output layer of the generator, we adopt the ReLU function, because the output represents gene expression that should be non-negative. To generate better performance, we utilize a structure similar to ResNet (He *et al.*, 2016), an identity skip connection, to construct our generator. To implement the ResNet structure, our layer is expressed in the following form:

$$
Y = T(BN(X, \{W_i\}) + X) \; T \in set(\text{Functions}), \tag{4}
$$

where $X$ represents the input data of our target layer, and $\{W_i\}$ represents a set of inner weight matrices. For the inner layers, $T = Mish()$, and for the output layer, $T = ReLU()$. From the perspective of back-propagation, the ResNet structure prevents gradients from being exactly zero, so the problem of gradient vanishing and white noise (very small gradient) can be solved to some extent (Balduzzi *et al.*, 2017). We use the skip connection strategy three times for each layer of decoder in our generator.

In most of the times, there are significant differences in gene expressions between different cell types, so theoretically, it is not difficult to distinguish batch-specific cell types (Cohen *et al.*, 2006; Ernst *et al.*, 2011). However, sometimes cells from different cell types may have similar gene expression levels. For such problem, we argue that the MNN method, our discriminator and adjusting training epoch could help keep a balance between mixing common cell types from different batches and preserving batch-specific cell types. Firstly, MNNs ensure that the pairs we select are the closest cells in each other's neighbors, so the gene expression levels are the most similar. Secondly, the discriminator is designed to discriminate cells from different batches. For batch-specific cell types, the discriminator should be able to distinguish them from cells in other batches quickly. Moreover, mixing these cells into other cell types can be treated as an over-fitting problem, so adjusting the number of training epochs and the size of mini-batches can help generate better results, depending on the chosen dataset.

In total, the training process of ResPAN is shown in Algorithm 2. The generator is trained once after training the discriminator 10 times and AdamW (Kingma and Ba, 2015) is used as our optimizer as recommended by the WGAN-GP paper. Tunable parameters for the neural network training are two coefficients $b_1$ (0.9) and $b_2$ (0.999) required by AdamW, number of epochs (300), batch size (1,024), learning rate (0.0001), and the value of $\lambda$ (10). Values shown in the parenthesis are the default settings.

#### 1.3 Metrics details

Details of all 16 metrics we used in the study are listed below. Note that for the two large-scale datasets (HCA and Mouse Brain), these metrics were calculated using 10% of randomly sampled data. This was repeated for 10 times and an average was taken over them to get the final metric for each combination of method and dataset.

- ASW: Average silhouette width (ASW), is computed based on the average intra-cluster distance and the average inter-cluster distance. The inter-cluster distance for a sample is calculated between it and its nearest cluster. More specifically, silhouette

**Algorithm 2** Training process with AdamW optimizer for AWGAN.

**Input:** Model $D$,$G$; Dataset $Data_{fake}$, $Data_{true}$; Mini-batch set $Batch$; Number of epochs $K$; Number of $D$ training before $G$ training $n_{critic}$.
**Output:** Model $G$; Dataset $Data_{integrated}$

INIT: initialize all parameters.
**for** $K$ steps **do**
    **for** $index, b$ in $enurmate(Batch)$ **do**
        $Refer = Data_{true}[b]$
        $Query = Data_{fake}[b]$
        $L_D = -E[D(Refer)]$
        $L_G = -E[D(G(Query))]$
        $x =$ Interpolation($Refer$,$G(Query)$)
        $Div = E[(\|\nabla_x D(x)\|_2 - 1)^2]$
        $L = L_D - L_G + \lambda Div$
        AdamW($D, L$) # Training D
        **if** $index$ % $n_{critic}$ == 0 **then**
            AdamW($G, L_G$) # Training G
$Data_{generated} = G(Data_{fake})$
$Data_{integrated} =$ concat($Data_{true}$,$Data_{generated}$)
Return $G, Data_{integrated}$

width is defined as:

$$SW = \frac{(d_{\text{inter}} - d_{\text{intra}})}{max(d_{\text{inter}}, d_{\text{intra}})}. \tag{5}$$

ASW is calculated by averaging across the silhouette width computed on each data point. Since we normally have two kinds of clusters: batch and cell type and we intend to remove batch effects and preserve different cell types, a model with low $bASW$ and high $cASW$ is desirable. To make positive direction indicate better performance, we use 1-$bASW$ instead of $bASW$.

- LISI: Local Inverse Simpson's Index (LISI) quantifies the degree of mixing in local neighborhoods. LISI chooses neighbors based on the local distribution of distances with a fixed perplexity and the chosen neighbors will be used to calculate the Inverse Simpson's Index. This metric was first used in Harmony (Korsunsky *et al.*, 2019). The definition of Inverse Simpson's Index is:

$$ISR = \frac{1}{\sum_{b=1}^{B} p(b)}, \tag{6}$$

where $b$ is the batch index (or cell type index) and $p(b)$ is the batch probability (or cell type probability) in the local neighborhood. There are two types of LISI: cell type LISI (cLISI) and batch LISI (bLISI). The former one measures the effective number of cell types in a local neighborhood, while the latter one measures the effective number of batches. An ideal mixing would result in cLISI being 1 and bLISI close to the number of batches. To make them bound between 0 and 1, we min-max scale them, that is subtracting 1 from them and then dividing them by either number of cell types -1 or number of batches -1. Our goal is to increase bLISI and decrease cLISI. Similar to ASW-based metrics, we use $1 - cLISI$ instead of $cLISI$ after the min-max scaling.

- Positive rate and TP rate: A cell is defined as positive if the percentage of its kNNs that have the same label as the cell is over 0.5, and positive rate is calculated as the proportion of positive cells in the corrected data. True positive (TP) rate is calculated only for positive cells, and a cell is TP if it is surrounded by appropriate proportions of cells from different batches. The appropriate proportions of cells can be determined based on the three sigma principle, that is, if we sample $k$ cells from cell type $n$, then the expected number of cells coming from batch $i$ should be $kp_i$ and the variance should be $kp_i(1-p_i)$, where $p_i$ is the proportion of cells from batch $i$ in all cells labeled as $n$. This idea was proposed in iMAP (Wang *et al.*, 2021).

- kBET: K nearest neighbor (kNN) batch-effect test (kBET) utilizes a kNN-based algorithm to measure batch effects (Büttner *et al.*, 2019). The algorithm randomly selects some data points and tests whether the local batch label distribution among them is similar to the global distribution. Ideally, the two distributions should be very close to each other, so the rejection rate should be close to 0. In fact, there are some differences between the two, so we get an actual rejection rate between 0 and 1. To be consistent with the direction of other metrics, we define the acceptance rate as:

$$\text{Acceptance Rate} = 1 - \text{Rejection Rate}. \tag{7}$$

  A higher acceptance rate of one model suggests its better performance on batch correction. For the neighborhood size ($k$), we set it as the average batch size times 0.25 and it is truncated between 10 and 100 if it is smaller than 10 or larger than 100.

- Graph Connectivity: This method is based on kNN and its target is to evaluate the mixing of common cell types after batch correction (Luecken *et al.*, 2021). The metric evaluates whether cells with the same cell type label are directly connected across batches. Let $T$ represents the set containing all cell type labels, we can use the following equation to compute the graph connectivity score:

$$gc = \frac{1}{|T|} \sum_{c \in T} \frac{|LCC\left(G\left(N_c, E_c\right)\right)|}{|N_c|}, \tag{8}$$

where $G(N_c, E_c)$ represents a kNN subgraph containing nodes from cell type c, $|LCC()|$ indicates the number of nodes in the largest connected component (LCC) of the given graph, and $|N_c|$ is the number of nodes with cell type label $c$. Based on the concept of LCC, a higher graph connectivity score suggests better batch correction.

- NMI: Normalized Mutual Information (NMI) is utilized to quantify the shared information between two clusterings. We utilize this metric to evaluate the performance of cell type label preservation by comparing Louvain clusters ($U$) computed on integrated data and the annotated cell clusters ($V$) from the same data. The metric is defined as:

$$NMI(U, V) = \frac{MI(U, V)}{GM(H(U), H(V))}, \tag{9}$$

where $H()$ is the entropy function and $GM()$ represents generalized mean. A larger NMI suggests better ability of a given method to preserve cell type labels. We calculated NMI in this paper using the scib Python package, where the Louvain clustering was optimized to obtain the best NMI for each dataset (Luecken *et al.*, 2021).

- ARI: Adjusted Rand Index (ARI) is also a metric for evaluating the overlap between two different clusterings. Still, we compare Louvain clusters obtained from integrated data and annotated cell type labels from the original data. Rand index takes into consideration both correct overlaps and correct disagreement between the two clusterings. The raw RI is adjusted for chance to get ARI and ARI is defined as:

$$ARI = \frac{RI - \text{Expected}(RI)}{max(RI) - \text{Expected}(RI)}. \tag{10}$$

A larger ARI indicates that a model has better performance on preserving cell type labels. We calculated ARI also using the scib Python package (Luecken *et al.*, 2021).

- kNN sim: K nearest neighbors (kNN) similarity is used to evaluate the preservation of local neighborhood around different cells before and after batch effect removal. An ideal batch correction method should preserve the information of local neighbors, which means that the neighbors of cells in one batch should remain largely unchanged before and after batch correction. For each batch, $k$ is set as 1% of the number of cells (truncated between 5 and 50) and this metric is calculated on the embedding space. For each batch, a kNN sim value is first calculated for each cell using Jaccard index. Then, an average score is obtained across all cells for the corresponding batch.

- Cell-cell sim: Cell-cell similarity preservation is used to evaluate the preservation of cell-cell similarity matrix before and after batch correction. Two cell-cell similarity matrices are calculated using cosine similarity on the embedding space of data before ($S_1$) and after batch correction ($S_2$) separately. The cell-cell sim is defined as:

$$\text{cell-cell sim} = exp\{-\frac{1}{N^2}\sum_{i,j}^{N}|S_{1ij} - S_{2ij}|\}, \tag{11}$$

where $N$ is the number of cells. In the paper, this metric was calculated in a batch-specific manner and the final cell-cell sim score was computed as the average across all batches.

- Exp sim: This is a metric for evaluating the conservation of biological information at the gene expression level. We believe that for cells within each batch, their expression profiles after removing batch effects should still be similar to the levels in the original data. Let $E_1$ and $E_2$ represent the $L_2$ normalized expression matrix for the original data and for the corrected data, respectively. The exp sim is defined as:

$$\text{exp sim} = exp\{-\frac{1}{N}\sum_{i}^{N}\sum_{j}^{G}(E_{1ij} * E_{2ij})\}, \quad (12)$$

where $N$ is the number of cells and $G$ is the number of HVGs. In our study, this metric was calculated first in each batch and an average value was then taken over all batches to get the final exp sim score.

- CC score: Cell cycle score is used to evaluate the preservation of cell cycle variations to the expression profile before and after batch correction. An ideal batch correction should retain the variance contribution of the S and G2/M phase scores. This score was calculated using the cell_cycle function from the scib package (Luecken *et al.*, 2021), which used the score_genes_cell_cycle function from the Scanpy package with a reference gene set provided by Tirosh et al. (Tirosh *et al.*, 2016) to compute the scores and used principal component regression to calculate the variance contribution before ($Var_{before}$) and after batch correction ($Var_{after}$) for each batch. The CC score is defined as :

$$\text{CC score} = 1 - \frac{|Var_{before} - Var_{after}|}{Var_{before}}. \quad (13)$$

- HVG score: This score is used to measure the level of HVG overlap between the data before and after batch correction. In addition, only methods that can complete batch correction on gene space can be used to calculate this score. Given that the

HVG set of data before batch correction is $G_1$, and that of data after batch correction is $G_2$, the HVG score is calculated as:

$$\text{HVG score} = \frac{|G_1 \cap G_2|}{min(|G_1|, |G_2|)}. \tag{14}$$

This score was calculated using the hvg_overlap function from the scib package (Luecken *et al.*, 2021).

- DEG F1: The F1 score of differentially expressed genes (DEGs) is used to evaluate the performance of different batch correction methods on preserving DEGs. We used the function rank_genes_groups from Scanpy to find two sets of DEGs per cell type in the data before batch correction $D_1$ and after batch correction $D_2$, then we treated $D_1$ as ground truth and computed the precision rate and recall rate. Note that for simulated data, since true DEGs were known, they were used as $D_1$. The value of DEG F1 is defined as:

$$\text{DEG F1} = \frac{2 * precision * recall}{precision + recall}. \tag{15}$$

##### 1.4 Device information

All analyses were performed on Yale's high performance computing clusters (Farnam). The CPU of our device is Intel(R) Xeon(R) Gold 6240, 2.6 GHz, and the GPU is NVIDIA TESLA A100, the RAM is 25GB.

#### References

Balduzzi, D. *et al.* (2017). The shattered gradients problem: If resnets are the answer, then what is the question? In *International Conference on Machine Learning*, pages 342–350. PMLR.

Büttner, M. *et al.* (2019). A test metric for assessing single-cell rna-seq batch correction. *Nature methods*, **16**(1), 43–49.

Cohen, S. M. *et al.* (2006). Denoising feedback loops by thresholding—a new role for micrornas. *Genes & development*, **20**(20), 2769–2772.

Ernst, J. *et al.* (2011). Mapping and analysis of chromatin state dynamics in nine human cell types. *Nature*, **473**(7345), 43–49.

Glorot, X. *et al.* (2011). Deep sparse rectifier neural networks. In *Proceedings of the fourteenth international conference on artificial intelligence and statistics*, pages 315–323. JMLR Workshop and Conference Proceedings.

Gulrajani, I. *et al.* (2017). Improved training of wasserstein gans. In I. Guyon, U. V. Luxburg, S. Bengio, H. Wallach, R. Fergus, S. Vishwanathan, and R. Garnett, editors, *Advances in Neural Information Processing Systems*, volume 30. Curran Associates, Inc.

Haghverdi, L. *et al.* (2018). Batch effects in single-cell rna-sequencing data are corrected by matching mutual nearest neighbors. *Nature biotechnology*, **36**(5), 421–427.

He, K. *et al.* (2016). Deep residual learning for image recognition. In *Proceedings of the IEEE conference on computer vision and pattern recognition*, pages 770–778.

Ioffe, S. and Szegedy, C. (2015). Batch normalization: Accelerating deep network training by reducing internal covariate shift. In *International conference on machine learning*, pages 448–456. PMLR.

Kingma, D. P. and Ba, J. (2015). Adam: A method for stochastic optimization. In *3rd International Conference on Learning Representations, ICLR 2015*.

Korsunsky, I. *et al.* (2019). Fast, sensitive and accurate integration of single-cell data with harmony. *Nature methods*, **16**(12), 1289–1296.

Luecken, M. D. *et al.* (2021). Benchmarking atlas-level data integration in single-cell genomics. *Nature Methods*, pages 1–10.

Misra, D. (2019). Mish: A self regularized non-monotonic neural activation function. *arXiv preprint arXiv:1908.08681*.

Satija, R. *et al.* (2015). Spatial reconstruction of single-cell gene expression data. *Nature biotechnology*, **33**(5), 495–502.

Tirosh, I. *et al.* (2016). Dissecting the multicellular ecosystem of metastatic melanoma by single-cell rna-seq. *Science*, **352**(6282), 189–196.

Tran, H. T. N. *et al.* (2020). A benchmark of batch-effect correction methods for single-cell rna sequencing data. *Genome biology*, **21**(1), 1–32.

Wang, D. *et al.* (2021). imap: integration of multiple single-cell datasets by adversarial paired transfer networks. *Genome biology*, **22**(1), 1–24.
