## Supplementary Figures for "ResPAN: a powerful batch correction model for scRNA-seq data through residual adversarial networks"

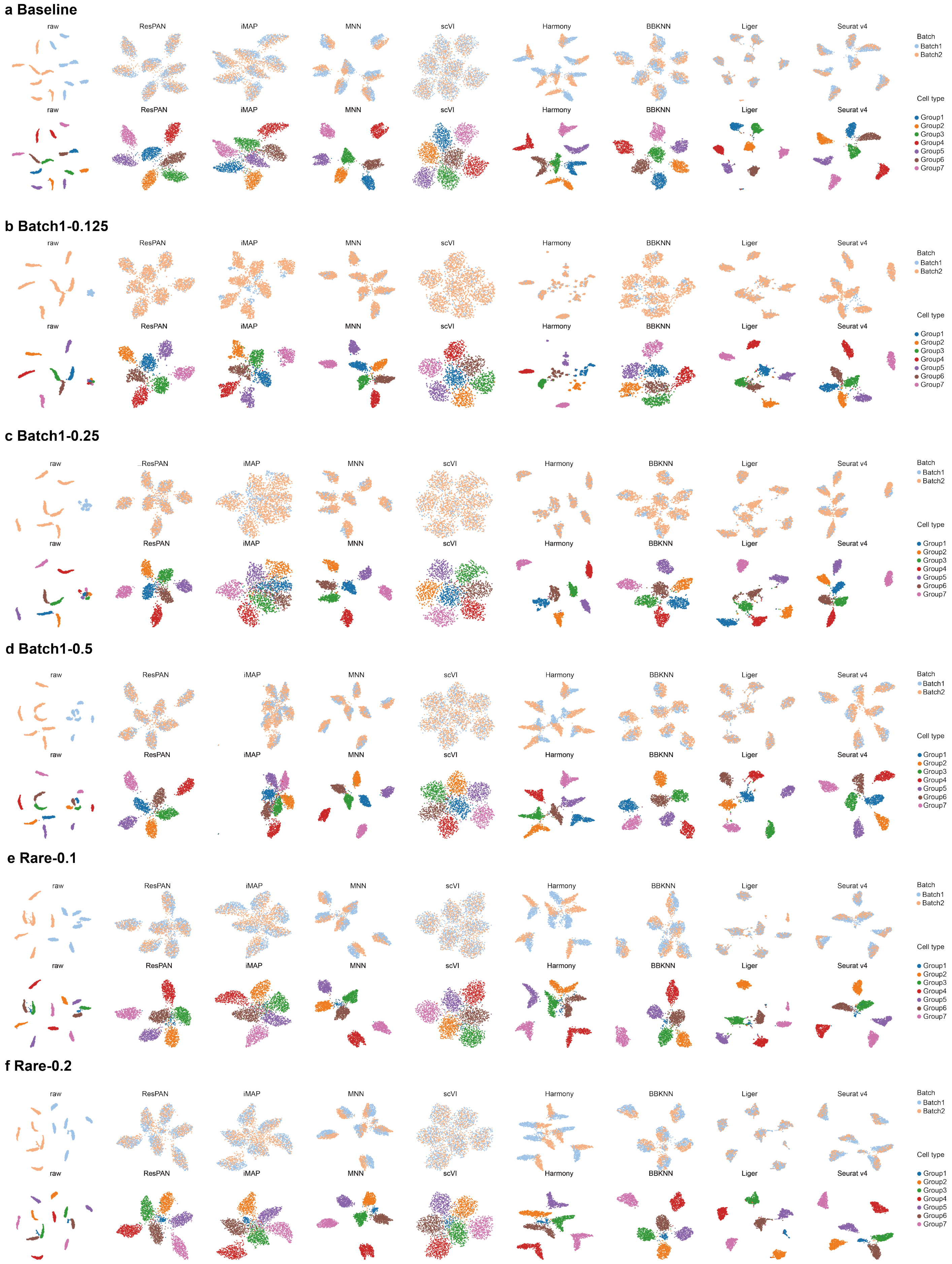
a Baseline
b Batch1-0.125
c Batch1-0.25
d Batch1-0.5
e Rare-0.1
f Rare-0.2
raw
ResPAN
iMAP
MNN
scVI
Harmony
BBKNN
Liger
Seurat v4
Batch
Batch1
Batch2
Cell type
Group1
Group2
Group3
Group4
Group5
Group6
Group7

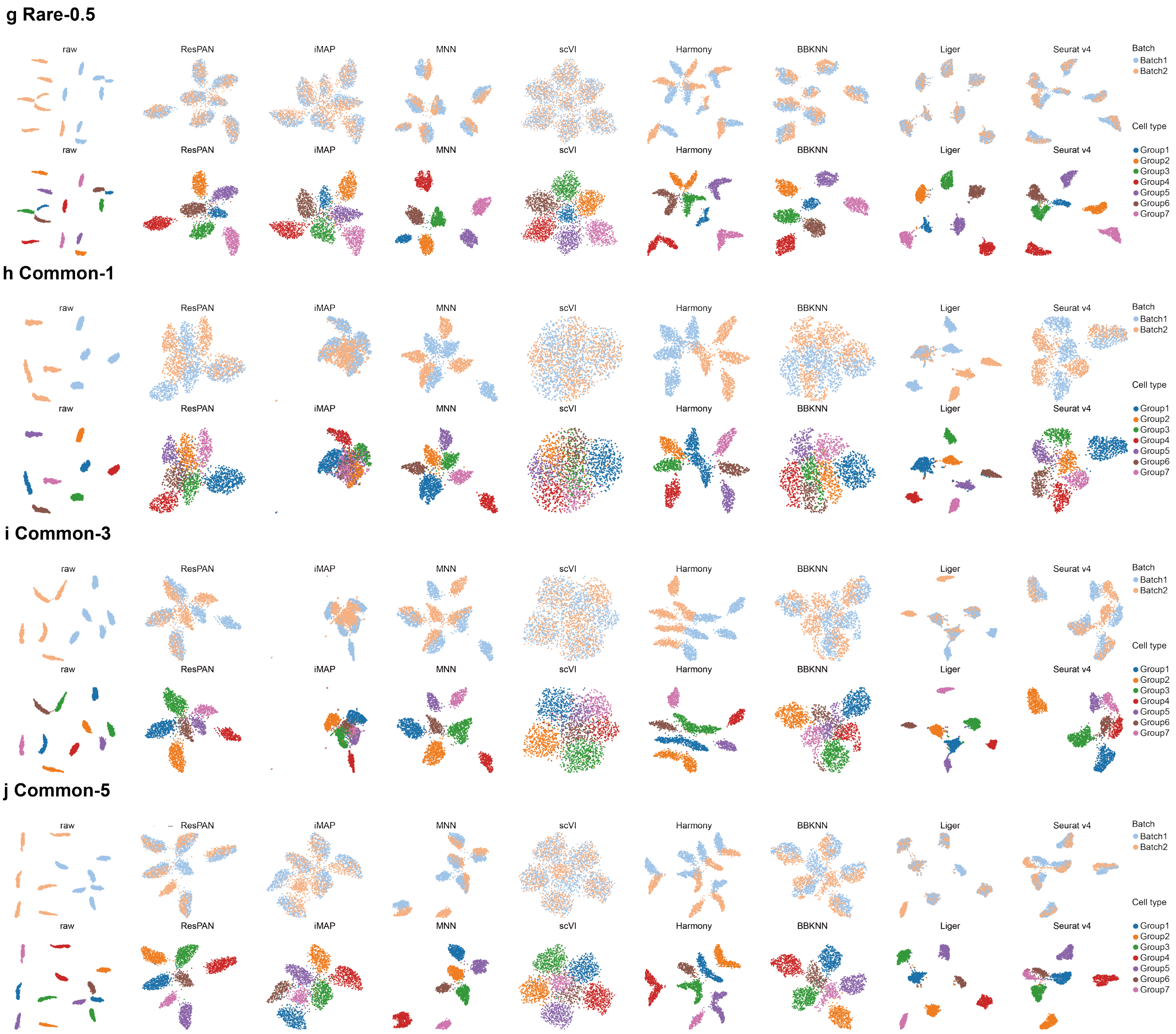


Figure S1: UMAP visualizations of simulated data under all ten sub-scenarios (seed 8). For each sub-figure, the first row are the UMAPs colored by batch labels, and the second row are the UMAPs colored by cell type labels.

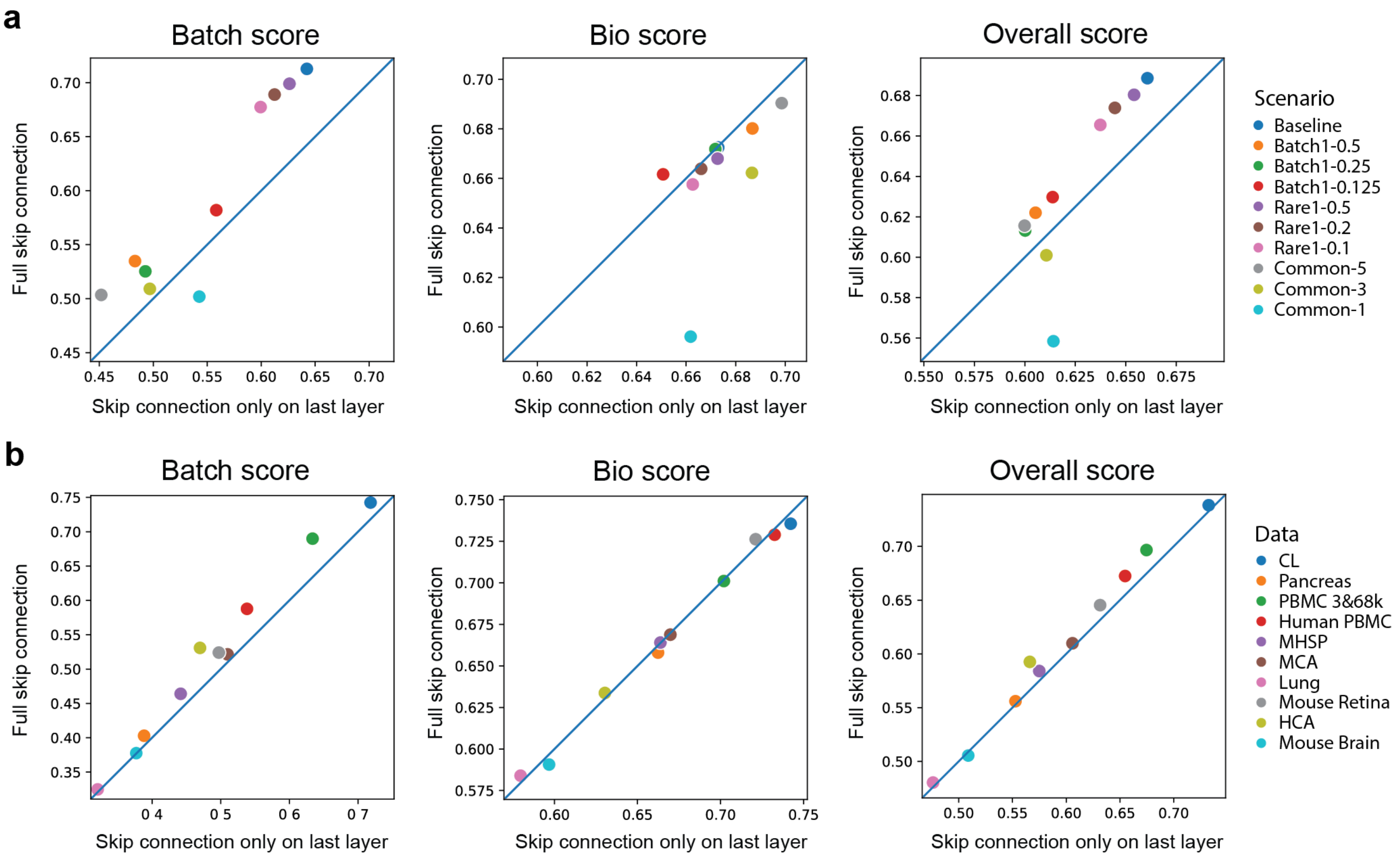


Figure S2: Performance comparison of full skip connection design and single skip connection design for the generator model on simulated data (a) and real data (b). Blue diagonal lines are lines with intercept at 0 and slope at 1, indicating the equal value of x and y axis.

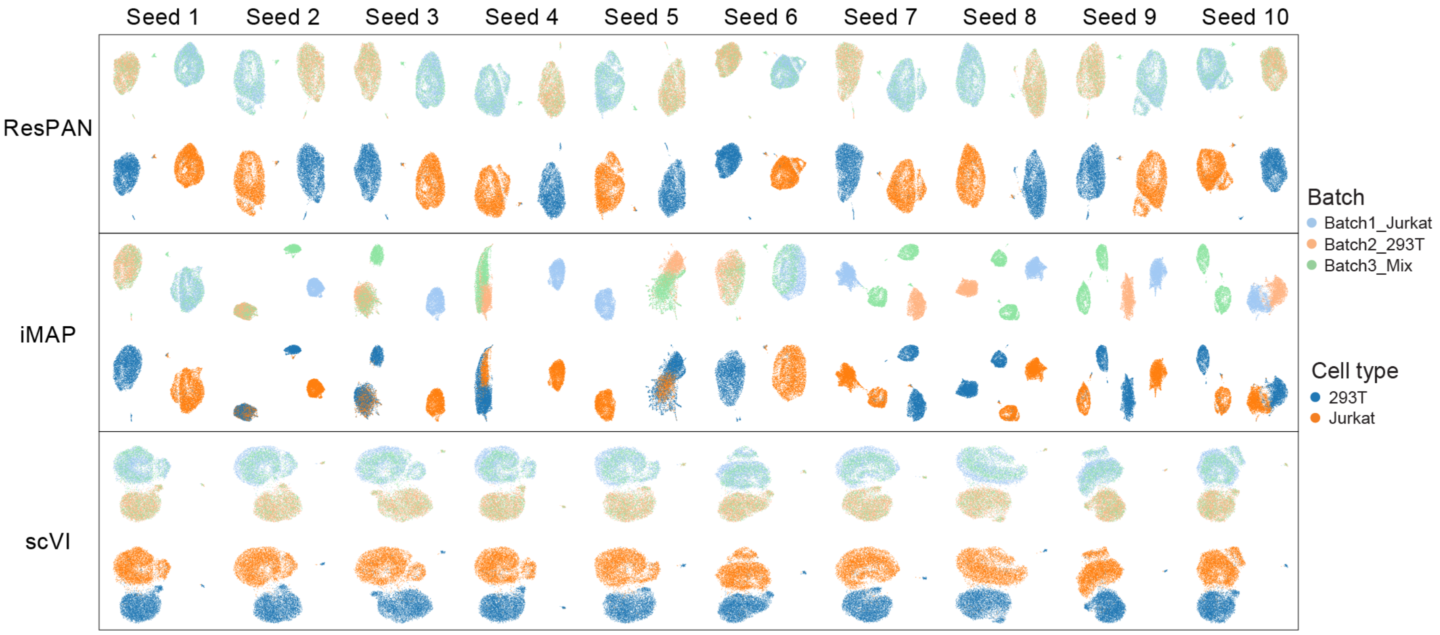


Figure S3: UMAP visualizations of the three deep learning methods (ResPAN, iMAP, and scVI) under ten seeds on the CL dataset. Each row represents a method and each column represents a seed.

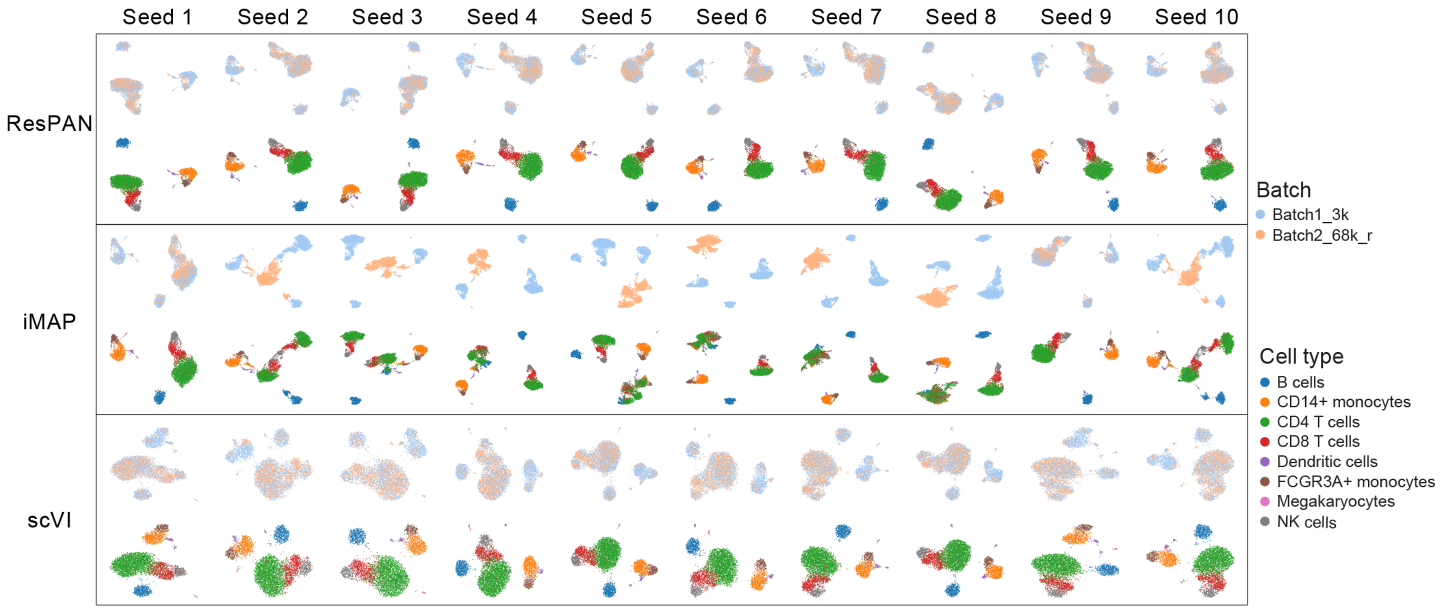


Figure S4: UMAP visualizations of the three deep learning methods (ResPAN, iMAP, and scVI) under ten seeds on the PBMC 3&68k dataset. Each row represents a method and each column represents a seed.

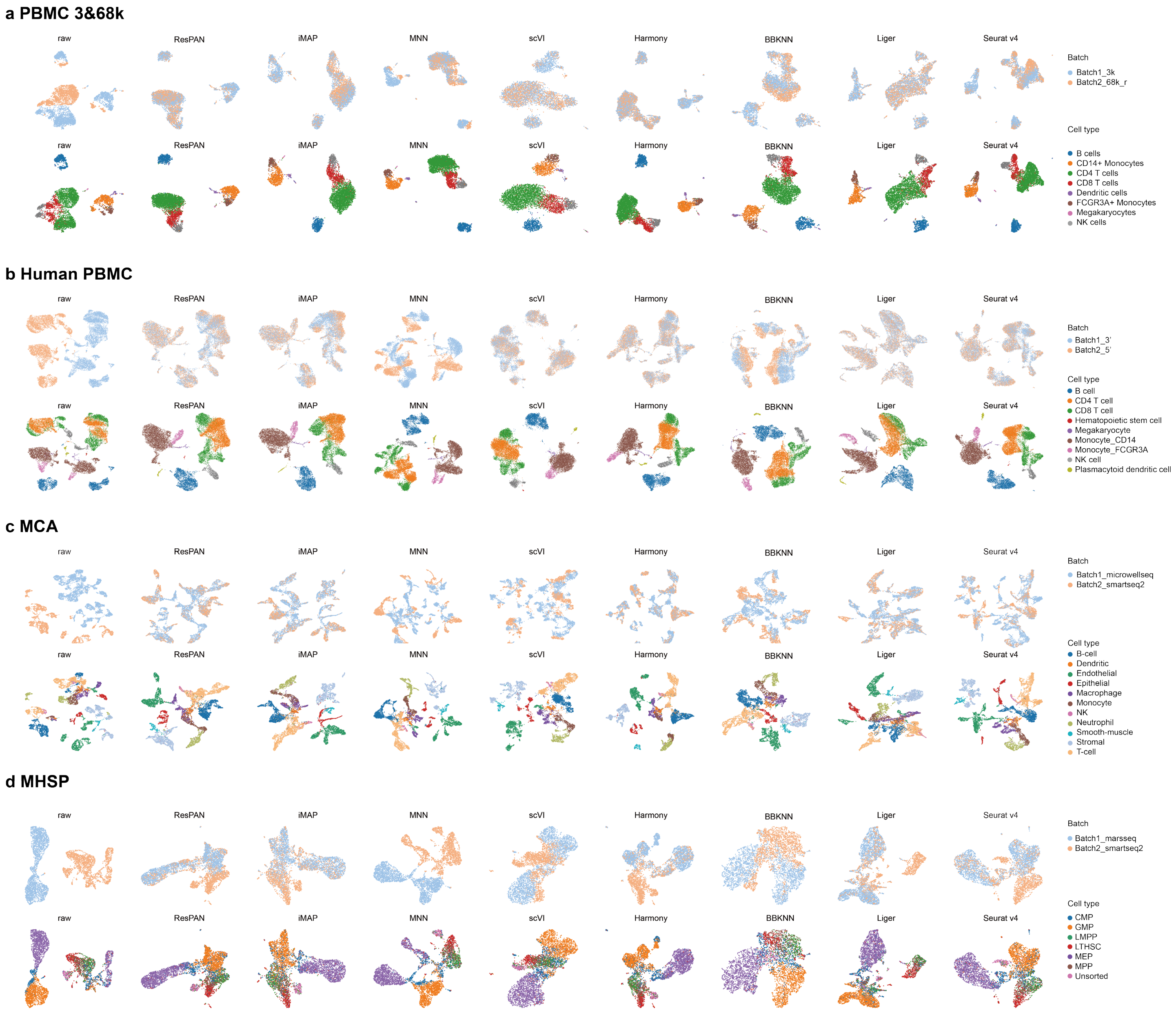

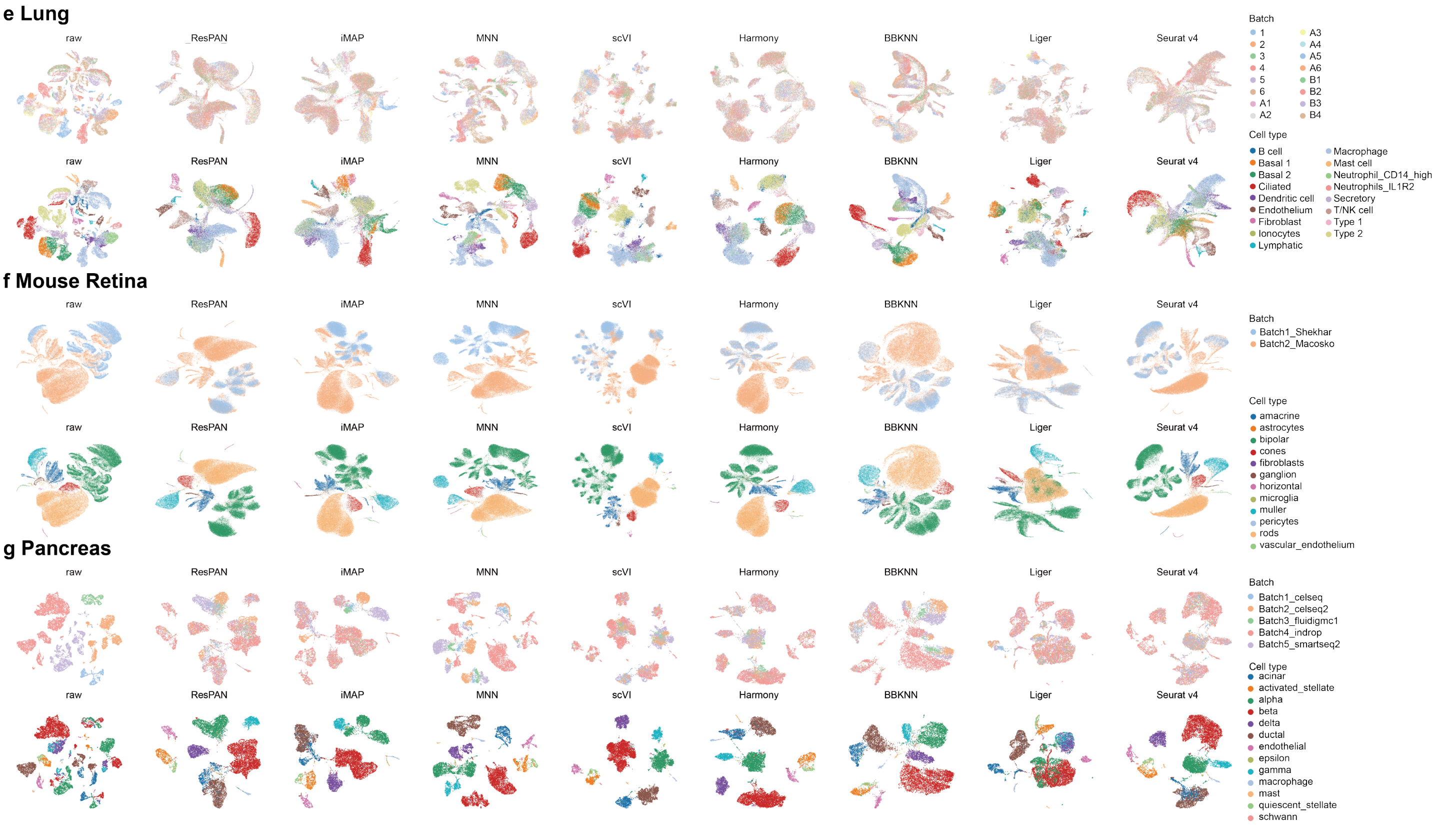


Figure S5: MAP visualizations of seven real datasets not shown in the main text. For each sub-figure, the first row are the UMAPs colored by batch labels, and the second row are the UMAPs colored by cell type labels.

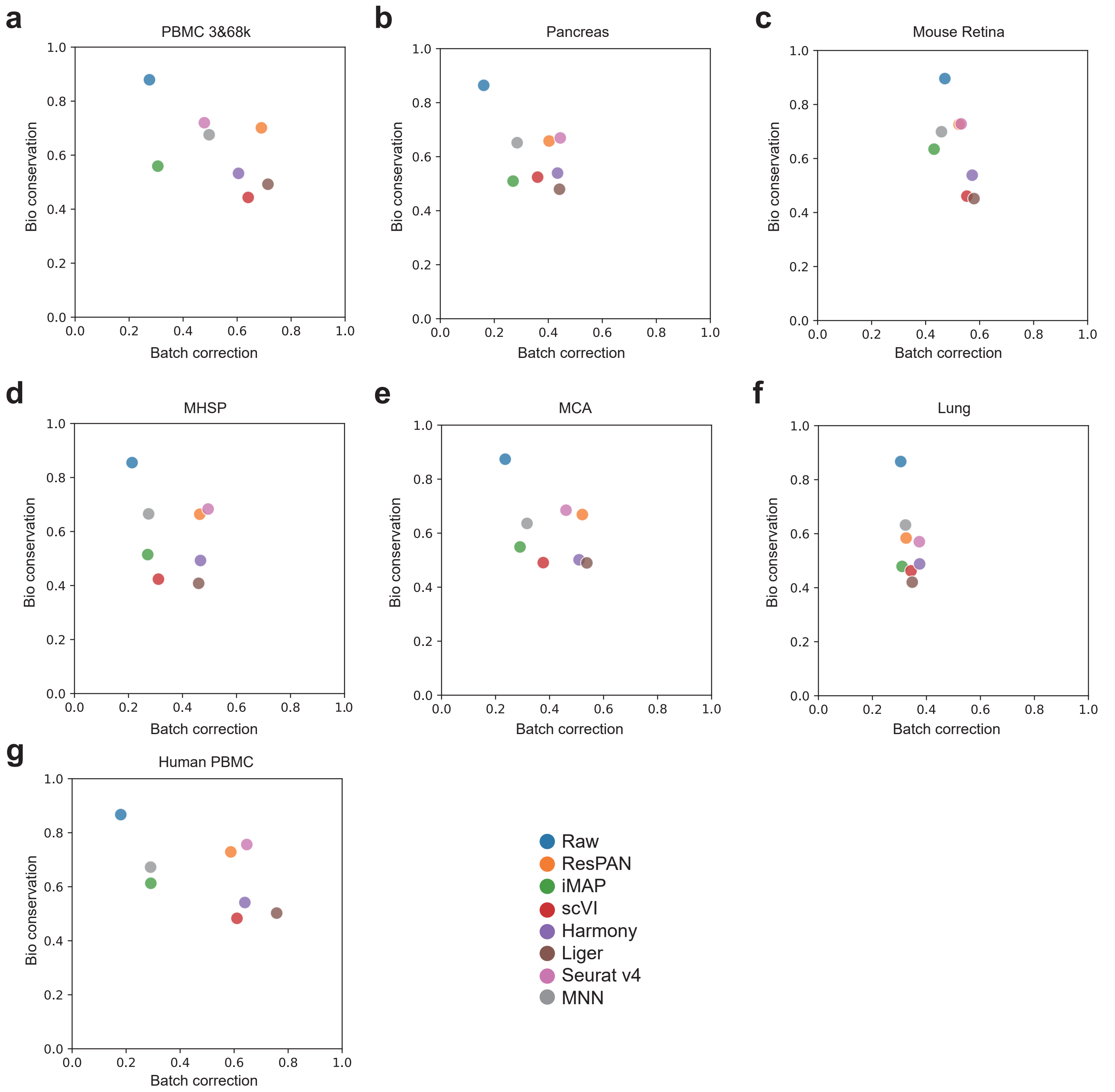


Figure S6: Bio conservation score versus batch correction score of seven real datasets not shown in the main text. For each sub-figure, the vertical axis represents the bio conservation score, and the horizontal axis represents the batch correction score.

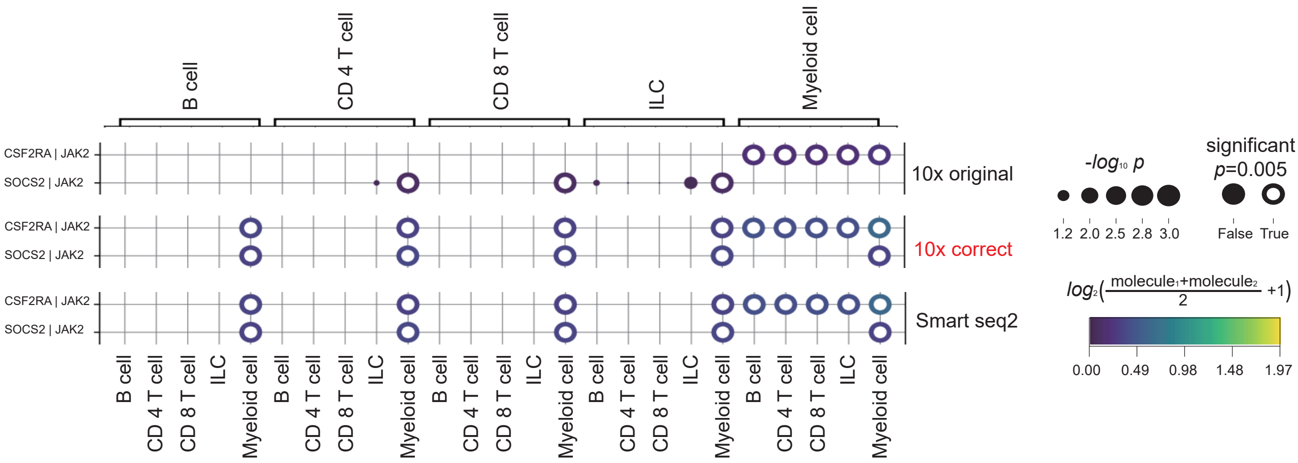


Figure S7: Novel pairs uncovered by ResPAN on the CRC dataset.

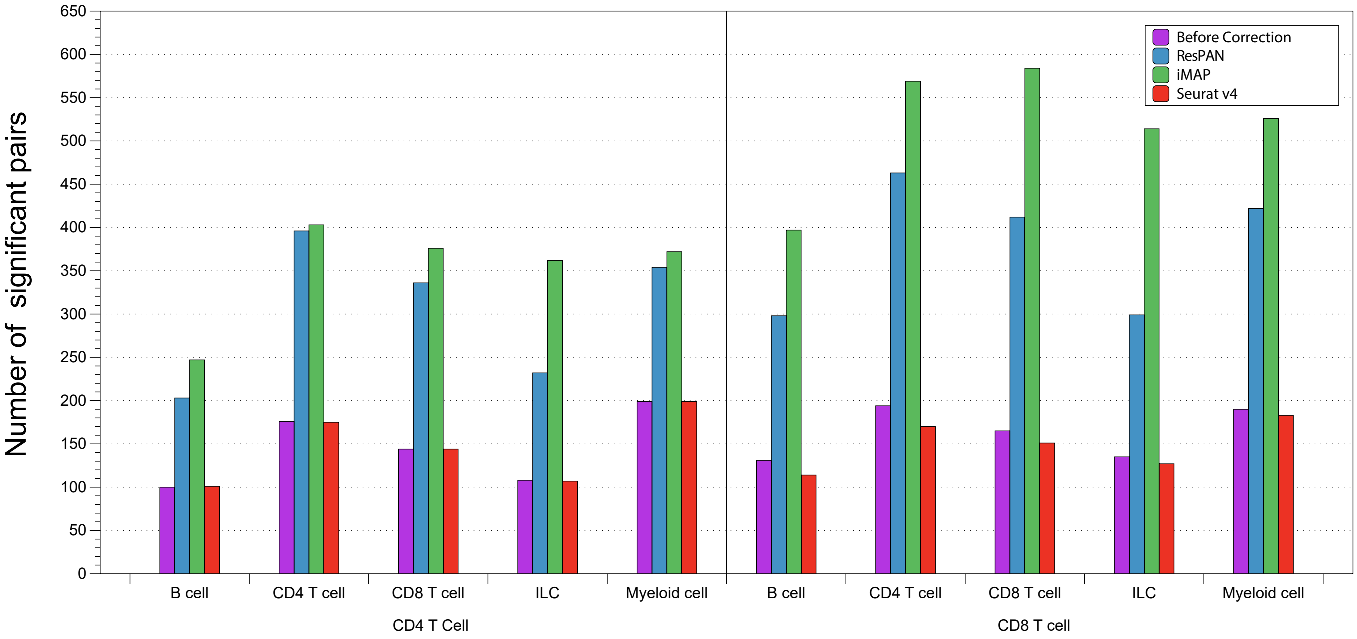


Figure S8: The number of significant ligand-receptor pairs form the CRC dataset generated based on raw data and three different batch correction methods.
